# Estimating fecundity and density dependence from mark-recapture data for making population projections

**DOI:** 10.1101/2020.08.26.268656

**Authors:** Bilgecan Şen, H. Reşit Akçakaya

## Abstract

Forecasting changes in size and distributions of populations is at the forefront of ecological sciences in the 21^st^ century. Such forecasts require robust estimators of fecundity, survival and density-dependence. While survival estimation is the main focus of mark-recapture modelling, fecundity and density dependence are rarely the subject of these models. Here, we demonstrate that these parameters can be simultaneously estimated in a Bayesian framework using only robust design mark-recapture data. Using simulated capture histories, we show that this framework (which we named CJS-pop) can estimate vital rates and their density dependence with little bias. When CJS-pop is applied to capture history data from Brown Creeper (*Certhia americana*), it provides estimates of fecundity that is expected from the breeding biology of this species. Finally, we illustrate that density dependence, even when estimated with uncertainty in the CJS-pop framework, regularizes population dynamics and reduces the frequent population extinctions and explosions observed under density-independent models. While CJS-pop as a whole is a useful addition to the current mark-recapture modelling toolbox, we argue that the independent components of this framework in estimating fecundity and density dependence can be integrated to other CJS frameworks, potentially creating models capable of population projections.

## Introduction

Mark-recapture data analysis is a staple in population ecology for estimating survival, abundance, and recruitment rates (Lebreton et al. 1992; Williams et al. 2002; Cooch and White 2016). More recently, mark-recapture methods have been extended to work in parallel with different types of data in frameworks such as integrated population models (IPMs; Schaub and Abadi 2011) and to estimate dispersal and animal movement in spatial capture-recapture analysis (Ergon and Gardner 2014; Schaub and Royle 2014). When considering the wide applicability of mark-recapture methods for estimating parameters related to population dynamics, rarity of two types of parameters in the mark-recapture literature stands out: 1) fecundity as defined in a population modelling setting; and 2) density dependence of survival and fecundity. If these parameters can be estimated from mark-recapture data, then this single data source can be used on its own to parameterize stage-structured population models.

Fecundity is a measure of breeding performance of a population and in the perspective of a PVA can be defined as per-capita number of offspring that survives to the next time step (Akçakaya et al. 1999). When no explicit data on fecundity are available, models such as Jolly-Seber and even some IPMs (for example, Ahrestani et al. 2017) estimate recruitment rate as the total number of individuals added to a population in a given time period (usually, immigrants and newborn individuals). Recruitment rate, in this regard, is not useful for making stage-structured population projections, because density dependence and stochasticity in fecundity cannot be explicitly accounted for in this parameter alone. A PVA that employs a stage-structured matrix model requires both the number of juveniles born per individual and the juvenile survival to be explicitly estimated (for example, Ryu et al., 2016).

Density dependence (DD) is the phenomenon of population vital rates being dependent on population size. Along with exogenous factors, it is one of the main natural processes that regulates population dynamics. There are examples of estimating the strength of density dependence of vital rates from mark-recapture data. For example, Ryu et al. (2016) provided a framework for estimation of fecundity and density dependence from robust design mark-recapture data. However, their framework is a mixture of frequentist and Bayesian approaches of mark-recapture models and it requires multiple models to be fit sequentially. As a result Ryu et al. (2016)’s different model components do not inform one another during estimation, which prevents making full use of the data at hand and propagating uncertainty in a hierarchical manner.

Here, we present an update on Ryu et al. (2016)’s framework. We use both simulated and real data to show that; 1) fecundity can be estimated alongside adult survival and capture probability in a single Bayesian framework using only robust design mark-recapture data; 2) estimating fecundity reduces uncertainty in juvenile survival estimates; and, 3) when these vital rates estimates are combined with density dependence, resulting stage-structured population models are useful for calculating conservation-relevant metrics.

## Material and Methods

### Model specifications

Two basic parameters are estimated in a standard CJS model (Cooch and White 2016): survival and capture probability. Robust design mark-recapture data separates these two processes to primary capture occasions (for example years), and secondary capture occasions within primary occasions (for example months). Populations are assumed to be closed (individuals don’t die or leave the population) among secondary occasions within a primary occasion. Capture probabilities are estimated for each secondary occasion, which can then be used to estimate the population size for a given primary occasion. Populations are assumed to be open among primary occasions, so individuals can leave the population or die. Survival is estimated with information coming across primary occasions. Below, for simplicity in presentation, we assume primary occasions are years and secondary occasions are months.

We denote *p_x,k,t,h_* as the monthly capture probability of a stage *x* individual at population *k*, year *t*, and month *h*; where, *x* = 1,2,3 …, *X*; *k* = 1,2,3…, *K*; *t* = 1,2,3…, *T*; and *h* = 1,2,3…, *H*. We can use this monthly capture probability to calculate a yearly capture probability:

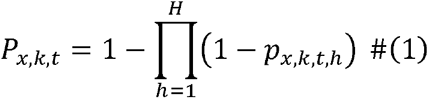

Then, we can use the heuristic estimator of populations size with a correction for years with no captures (Dail and Madsen 2011) to estimate the expected abundance of stage *x* individuals at population *k*, year *t*:

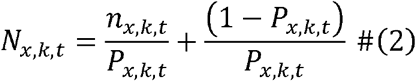

Where, *n_x,k,t_* is the number of captured stage *x* individuals in population *k* and year *t*. Using the expected abundance time series of each population we calculate a density index:

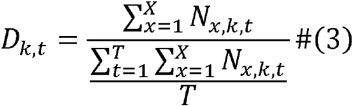

where, *D_k,t_* is the density index at population *k*, year *t*. The numerator in equation 3 is the total abundance of population *k* at year *t* across all stages, and denominator is the average expected total abundance across T years. *D* is an index for the deviation of population abundance in a given year from the long term average population abundance and it can be considered as relative population density. We use *D* as a covariate for estimating density dependence strength of fecundity and survival.

All three parameters estimated by equations 1 to 3, which are yearly capture probability, expected abundance, and density index, respectively, can be used to estimate fecundity and its density dependence:

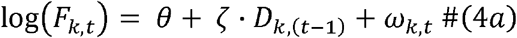

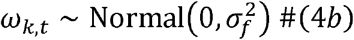

where, *θ* is the fecundity in log scale at 0 density; *ζ* is the change in fecundity in log scale with one unit change in population density index; *ω_k t_* is the spatio-temporal random effect at population *k* and time *t*; and 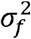 is the spatio-temporal variance of fecundity at log scale. We link the fecundity estimate to the number of captured juveniles in a given year and population by using expected abundances for adults calculated in equation 2. Below we present a simple case for two stages where *x* = 1 are juveniles and *x* = 2 are adults:

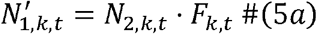

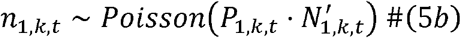

The expected number of juveniles is estimated twice in this framework: once as a derived variable (*N*_1,*k,t*_) using the heuristic population size estimator, and a second time 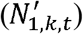 as a function of density index and expected number of adults. We discuss this double estimation and our choice of Poisson distribution in detail in discussion section below.

So far, we have only used captured number of juveniles and adults as a data source. However, this form of a capture history is not enough to estimate parameters used in the above 5 equations. So, we link these 5 equations with a more typical CJS model where survival and capture probability are used to model capture histories of individuals:

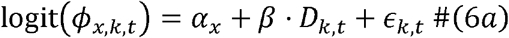

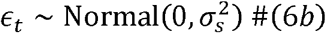

where, *ϕ_x,k,t_* is the apparent survival probability of a stage *x* individual at population *k* and year *t*; *α_x_* is the survival probability of a stage *x* individual on logit scale at 0 density; *β* is the change in survival in logit scale with one unit change in population density index; *ϵ_k,t_* is the spatiotemporal random effect at population *k* and time *t*; and 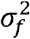 is the spatio-temporal variance of survival at logit scale.

Apparent survival changes the latent states of individuals in a population between time steps, from t to *t* + 1; this latent state indicates whether an individual is alive and in the population (*Z* = 1), or it is dead or left the population (*Z* = 0). The latent state of the *i*th individual, then, is determined by its state at time *t* and its survival to time *t* + 1.

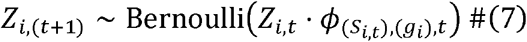

where, *ϕ*_(*s_i,t_*),(*g_i_*),*t*_ is the apparent survival probability at the breeding stage and population of the ith individual from year t to *t* + 1; *S_i,t_* is a matrix indicating the breeding stage of the *i*th individual in year *t*, and the vector *g_i_* indicates the population that the *i*th individual is in. Latent state, *Z*, also determines the potential capture of an individual; dead ones cannot be captured.

Hence, every element of the capture history, *y_i,t,h_* (1 if an individual is captured, 0 if it is not), is a Bernoulli random variable with a monthly capture probability, *P*_(*s_i,t_*),(*g_i_*),*t,h*_, conditional on the individual being alive and in the population, *Z_i,t_*.

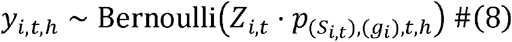

For simplicity in referring to this framework, we named it CJS-pop; pop extension comes from the fact that it can estimate necessary parameters to build a stage-based population model: Fecundity, staged-based survival, density dependence, and process variance.

### Simulated data

We simulated several sets of capture histories in order to test CJS-pop’s ability to correctly retrieve true parameter values, and to uncover any inherent biases, especially when quantifying density dependence strength. We set up a simulation scheme where we explored the effect of sample size on quantifying density dependence strength. We set the time series length to 17 years, which is the maximum time series length for Brown Creeper data set we are using (see below). We simulated three cases with 1, 5, and 10 populations, and three carrying capacities (which controls population size): 50, 100, 150. For each combination of the number of populations and carrying capacity, we generated capture history data sets using weak, moderate and strong density dependence on survival and fecundity, which created 27 separate simulation sets. For each simulation set, we generated 56 capture history data sets and fitted CJS-pop to each one. See Appendix S2 for more detailed discussion of the simulation framework and Appendix S3 for its code.

### Empirical data: Brown Creeper

We applied CJS-pop to Brown Creeper (*Certhia americana*) data obtained from the Mapping Avian Productivity and Survivorship (MAPS) program. Brown Creeper is a widespread North American songbird species. We treated each MAPS location (a cluster of mist-netting and banding stations) of Brown Creeper to be a separate population. We only included data from populations which were located in the contiguous U.S., and which have been monitored for at least 5 years. This resulted in a data set with 2931 individuals. We categorized any individual in its first year as a juvenile (MAPS age codes 2 and 4), and older individuals as adults (MAPS age codes 1, 5, 6, 7, and 8).

We made several adjustments and additions to basic CJS-pop framework presented above when applying to Brown Creeper data. First, we accounted for potentially transient individuals in the data set. In a CJS model, estimated survival rates are said to be “apparent” because the estimated survival rate cannot distinguish between dead individuals and the ones that just left the population. This can bias survival estimates to be lower than their true values. Accounting for transients is a partial way to correct for this bias and it is a frequently used technique in CJS literature (for example, Ahrestani et al. 2017).

Second, we used the priors and population modelling structure of CJS-pop to our advantage to estimate a juvenile survival rate with less uncertainty. We use information from adult survival and fecundity estimates, and the fact that they are density dependent, to estimate juvenile survival:

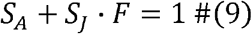

where, *S_A_* and *S_j_* are survival rates of adults and juveniles at mean population size (when *D* = 1) and *F* is the fecundity at mean population size. This equation states that population growth rate (*λ*) is equal to 1 when population abundance is at its long-term average. Using this equation, we would only need priors for adult survival and fecundity, and we can calculate juvenile survival as:

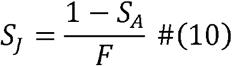

Third, we used a zero-inflated Poisson for modelling fecundity in equation 5b because there were several years and populations with no juvenile captures. Fourth, we did not use populations and years with no adult captures when modelling fecundity. While we account for no capture years in equation 2, we believe that limiting fecundity estimation to years with adult captures provides a more robust estimate.

Fifth, we changed the spatio-temporal variance structure of survival to be only temporal. Spatio-temporal variance structure in survival for Brown Creeper proved to be problematic because it made convergence harder for multiple parameters while also reducing the effective sample size of their MCMC chains. Lastly, we standardized the density index (*D*), calculated in equation 3, to be 0 at mean population size. This allowed for faster convergence of the MCMC chains in JAGS. See Appendix S2 for details on these adjustments to basic CJS-pop framework and goodness-of-fit testing of CJS-pop.

We fit three different CJS-pop models to Brown Creeper data: 1) density dependent, 2) density independent, and 3) Density dependent without the residency model.

### Population Projections

We ran population projections using a stage-structured population model with environmental and demographic stochasticity in both survival and fecundity. We parameterized these population models with 3 different parameter sets:

1. True simulation parameters that we used to generate capture history data.
2. Parameter estimates from density-dependent and density-independent CJS-pop fit to simulated data. At each replication we randomly selected the parameters of CJS-pop fit to one of the 56 data sets with this population and carrying capacity combination. To further incorporate parameter uncertainty, we randomly used at each replication the 2.5%, 50%, or 97.5% percentiles of the selected parameters.
3. Parameter estimates obtained from CJS-pop fit to the Brown Creeper data. We employed the full posterior distribution of the estimated parameters. At each iteration of the projections, parameter estimates were randomly selected from the posterior distributions of all parameters with respect to their correlation structure.

Using each of these sets of parameters, we ran single-population projections, with a carrying capacity of 1000 and an initial population of 500 adults and 500 juveniles. We ran the projections with 1000 replicates, and each replicate for 20 years. In order to incorporate environmental stochasticity, at each iteration and at each year we generated random temporal effects separately for survival and fecundity using 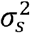 and 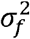, respectively. We recorded the minimum abundance of the population across 20 years for each iteration, and the distribution of minimum abundance among iterations. The expected value of this distribution is called expected minimum abundance (EMA) and it is a more informative metric than extinction risk, because the latter often has a distribution restricted to near-zero or near-one values (McCarthy and Thompson 2001).

### Software

See appendix S2 for a list of R packages used in the analysis. We used JAGS (Plummer 2003) as the MCMC sampler when fitting CJS-pop to data. We ran the models with 4 chains, 50000 iterations, 20000 burn in, and a thinning rate of 10 for simulated data sets, and with 4 chains, 100000 iterations, 50000 burn in and a thinning rate of 20 for Brown Creeper data. We checked convergence with R-hat values, and assumed chains were converged when 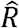 was <1.05. See Appendix S4 for JAGS code of CJS-pop. Additionally, R and JAGS code of CJS-pop analysis, example simulation data, and results of Brown Creeper analysis are accessible as Data S1 and also at https://github.com/bilgecansen/CJS-pop. The source code and data has been archived and made accessible in Zenodo (DOI: 10.5281/zenodo.3736702).

## Results

### Simulations

CJS-pop is able retrieve true parameter estimates without any apparent bias except for density dependence (Appendix S1: Figs. S1-6). DD strength is estimated with no bias when strength of DD used to generate capture history data is moderate. Strong DD in data simulation leads to slight underestimation of DD strength. There is, however, considerable overestimation of true DD parameters when capture history simulation was carried out with weak DD strength (Appendix S1: Fig. S1).

### Empirical Example: Brown Creeper

We detected weak DD on survival (*β* = −0.27, Fig. 1a), and on fecundity (*ζ* = −0.13, Fig. 1b) for Brown Creeper. Process variance estimations are low for survival, and high for fecundity, (*σ_s_* = 0.23, *σ_f_* = 0.97; Table S1). Survival at mean population size for adults and juveniles were estimated at 0.42 and 0.31, respectively. Our estimate of fecundity at mean population size was 1.94 (1.21 – 3.01) juveniles per adult (Appendix S1: Table S1). In addition, Bayesian p-values for both the survival and fecundity components indicated good model fits (0.24 and 0.50, respectively).

**Figure 1:**
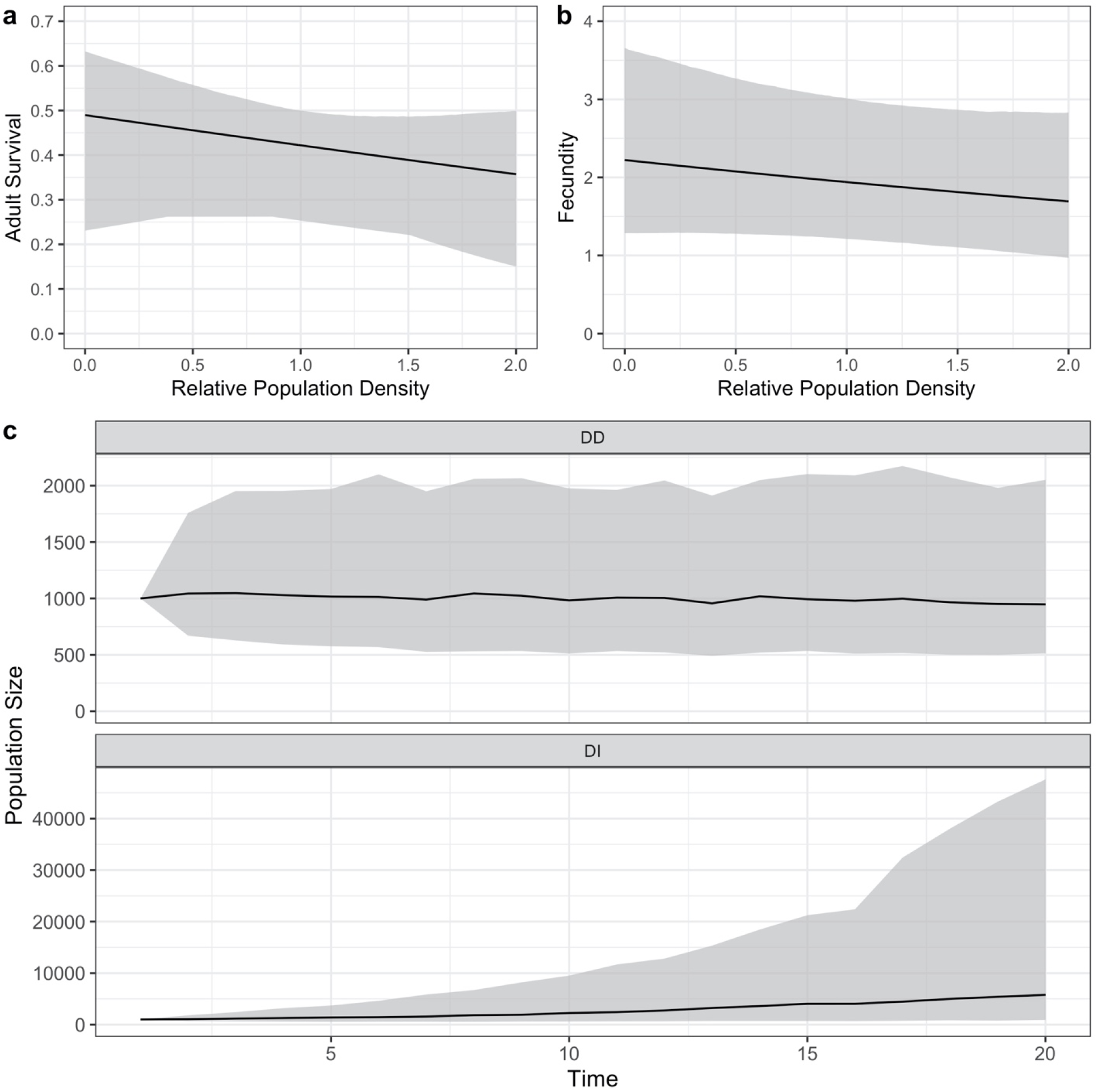
a) Relationship between adult survival and relative population density (*D*) of Brown Creeper as modeled by CJS-pop. b) Relationship between fecundity and relative population density (*D*) of Brown Creeper as modeled by CJS-pop. c) Solid line indicates the median trajectory of population size, across 12000 trajectories, of Brown Creeper projected by a stage-structured population model that was parameterized with a Density-Dependent (DD) or Density-Independent (DI) CJS pop. Shaded areas include 50% of the population trajectories. Carrying capacity was set to 1000 in the DD projections. Note the difference in scale in population size between DD and DI projections.

### Projections

Density-independent projections parameterized with CJS-pop fit to Brown Creeper data lead to frequent population extinctions and explosions, which is apparent in the population trajectory (Fig. 1c) and bi-modal distribution of minimum abundances (Fig. 2). Density dependence in population models lead to more regularized projections in which population extinctions and explosions are less frequent (Fig. 1c and Fig. 2). The distribution of minimum abundances from density-dependent projections of Brown Creeper show a generally similar pattern to projections parameterized with CJS-pop fit to simulated data irrespective of the DD strength of capture history simulations (Fig. 2).

**Figure 2:**
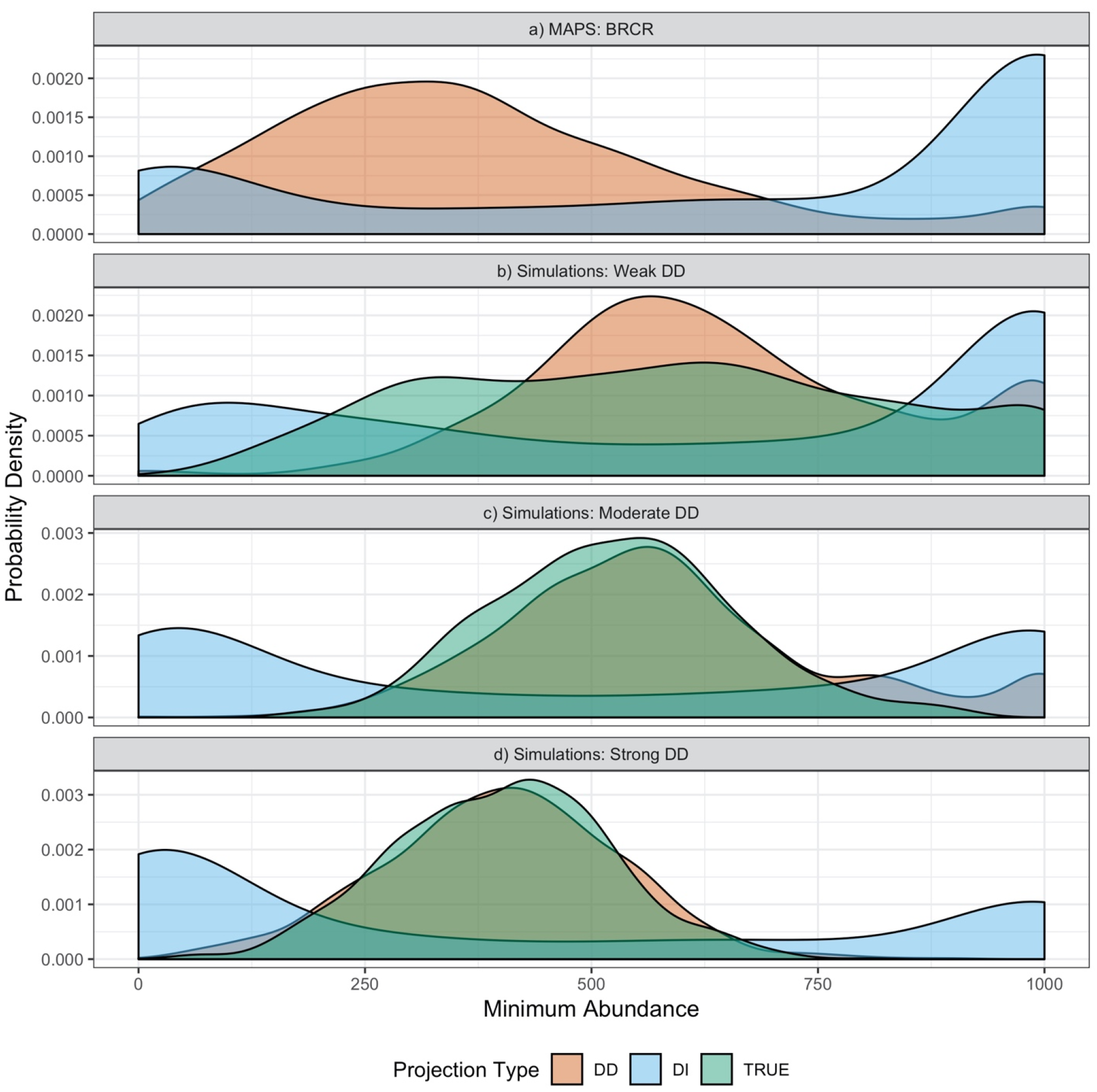
Distributions of minimum abundances resulting from population projections made with stage-structured population models that were parameterized with CJS-pop fit to Brown Creeper data (a), and with CJS-pop fit to simulation sets that was generated with different density-dependence strengths (b,c,d). Light blue represents parameterizations of population models with density-independent (DI) CJS-pop, orange represents density-dependent (DD) CJS-pop, and green (TRUE) represents population models that was parameterized with original simulation parameters that was used to generate capture history data. High probability density at 0 and 1000 indicates frequent population extinctions and explosions, respectively.

Projections made with population models that are parameterized with density-dependent CJS-pop fit to simulated data are close to projections made with true simulation parameters, especially when true simulation parameters included moderate or strong DD. This correspondence demonstrates, in a biologically relevant context, the ability of CJS-pop to fit realistic models to data (Figs, 2c-d). In contrast, projections with density-independent CJS-pop (Figs. 2b-d), and with CJS-pop fit to simulated capture history data with weak DD (Fig. 2b) were not close to projections with the true simulation models. Overestimation of DD strength in CJS-pop fit to weak DD data also results in overestimation of projected minimum abundances. Density-independent projections tends to result in frequent population extinctions or explosions (Figs. 2b-d).

## Discussion

CJS-pop is ready to be applied to bird species captured in the MAPS program for making population projetions. It can be extended to include weather, climate and other exogenous factors in addition to population density. The true value of CJS-pop lies in its ability to use limited data to parameterize population models that in turn can be used to predict changes in population sizes and distributions. We believe this is especially important for developing countries that do not yet have extensive bird banding and survey programs like MAPS and BBS. We don’t, however, think that whole CJS-pop framework needs to be used for the work we presented here to be useful. Rather, we argue that the independent ideas explained in estimating fecundity, juvenile survival and density dependence can be integrated to other CJS frameworks, potentially creating models capable of population projections. Below we describe 4 of the main advances CJS-pop provides to mark-recapture literature and discuss the trade-offs made when building the framework.

### 1 Fecundity Estimation

We estimated fecundity in the simulation data with no apparent bias (Appendix S1: Fig. S4). The fecundity estimate for Brown Creeper (*F* = 1.94 (1.21 − 3.01); Table S1) is also biologically realistic; Brown Creepers tend to have a single brood with a clutch size of 5 to 6 eggs in a breeding season. The main contribution here is that every vital rate (including fecundity), capture probabilities, and nuisance parameters were estimated simultaneously in a single model run. This allows for propagation of uncertainty among these parameters but also makes it possible for parameters that are estimated with less data (juvenile survival) to be informed by parameters estimated with more data (fecundity and adult survival).

### 2 Juvenile Survival Estimation

We detected no biases in juvenile survival estimates from CJS-pop in the simulation data (Appendix S1: Fig. S2). Juvenile survival estimates of Brown Creeper were similar between a density-dependent CJS-pop, and a CJS model that did not include fecundity or density dependence estimation but accounted for “transient” juveniles that leave their population in their first year (0.32 and 0.30, respectively). However, using information from fecundity and adult survival in setting the prior for juvenile survival reduced the estimation uncertainty considerably in CJS-pop. The 95% credible interval for juvenile survival in the CJS model is 0.08 − 0.71, while in CJS-pop this interval is 0.19 − 0.48.

### 3 Density Dependence Strength Estimation

Density dependence strength in mark-recapture studies is usually estimated using abundance directly as a covariate (for example, Nater et al. 2018). This approach can become problematic with more than one population, especially when each population has different habitat characteristics and therefore can support different number of individuals. If the goal is to estimate a species-specific density dependence strength that is applicable across all populations of the species, abundance of each population in each time step should be standardized with a proxy for how many individuals each population can support (e.g. carrying capacity). Here, we made this standardization using long-term abundance average of each population (equation 3). The density dependence strength we estimate is minimally biased for capture histories generated by moderate and strong density dependence (Appendix S1: Fig. S1). There is a more apparent bias when capture histories are generated with weak density dependence (Appendix S1: Fig. S1). However, the weak density dependence strength we used does not constitute a biologically realistic scenario if we consider the intrinsic growth rate associated with the density dependence strength from an allometric standpoint (See Appendix S2 for a detailed discussion about parameterizing simulations). Most importantly, density dependence strength estimates of survival and fecundity are ready to be used for population projections.

### 4 Population Projections

Estimation of fecundity, juvenile survival, and density dependence strength makes it possible to make population projections that are useful for conservation purposes. All of these three vital rates and demographic parameters are required, in addition to what is estimated in standard CJS models, to build a stage-structured population model. However, instead of fecundity, a parameter called recruitment rate is frequently estimated from frameworks such as IPM (for example, Ahrestani et al. 2017). This parameter cannot be used in stage-structured population models because it combines information from both emigration rate and the number of new-born individuals. Additionally, including only fecundity estimation in a CJS framework is also not enough, because without density dependence, these population projections would not be biologically meaningful. Population projections tend to explode in size or go extinct under exponential growth when there is environmental stochasticity that is not regularized by density dependence (Fig 1). The usefulness of this regularizing effect is also visible in the distribution of minimum abundances from population projections. A density-independent model cannot capture the minimum abundance distribution generated with a stochastic and density-dependent simulation, even when density dependence is weak. (Fig 2).

### Trade-offs

Using robust design mark-recapture data as the sole data set for the CJS-pop framework requires several trade-offs. First, we estimate expected juvenile abundance twice (*N*_1_ in equation 2 and 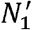 in equation 5a). If we only use 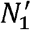, JAGS will give an error regarding the circular structure of the model because 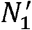 would also have been used in the denominator of equation 3. We see this as a minor issue because 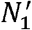 is used in the estimation of fecundity and we showed that fecundity can be estimated without bias in this structure (Appendix S1: Fig. S4).

Second, we estimate population sizes as expected values rather than random variables in CJS-pop framework. Modelling population sizes as random variables either requires informative priors on population sizes themselves or another data set, such as population counts, to make the model more stable and allow convergence (this is essentially what IPMs do). Here, however, we wanted to show that the vital rates necessary for population models can be estimated using only mark-recapture data. Using expected abundance, while not ideal, ensures that this framework requires only a single data source.

Third, we use a Poisson distribution instead of a binomial in equation 5b. Number of captured juveniles cannot be higher than the actual number of juveniles; this relationship is explicitly modelled as such with a binomial distribution. However, because we are using expected abundances in CJS-pop, there could be instances when there are more captured individuals than the expected abundance. Poisson distribution, by allowing such instances to occur, increases model stability and eases convergence. Because Bayesian p-value for brown creeper data is 0.5, we can say that this structure can represent the data well (Bayesian p-values close to 0.5 indicate better fit; Kéry and Royle 2016). Last but not least, the framework we present here is complex and in JAGS it takes for about 20 hours for Brown Creeper model to converge.

## Acknowledgements

We thank Kevin Shoemaker for valuable input. This study was initiated under funding from the NASA Biodiversity Program (NNH10ZDA001N-BIOCLIM). We thank the many volunteers who have contributed to the MAPS program and the Institute for Bird Populations for development and active curation of the MAPS dataset.

