## Appendix S1 for "Estimating fecundity and density dependence from mark-recapture data for making population projections"

### Appendix S1: Supplementary Figures and Tables

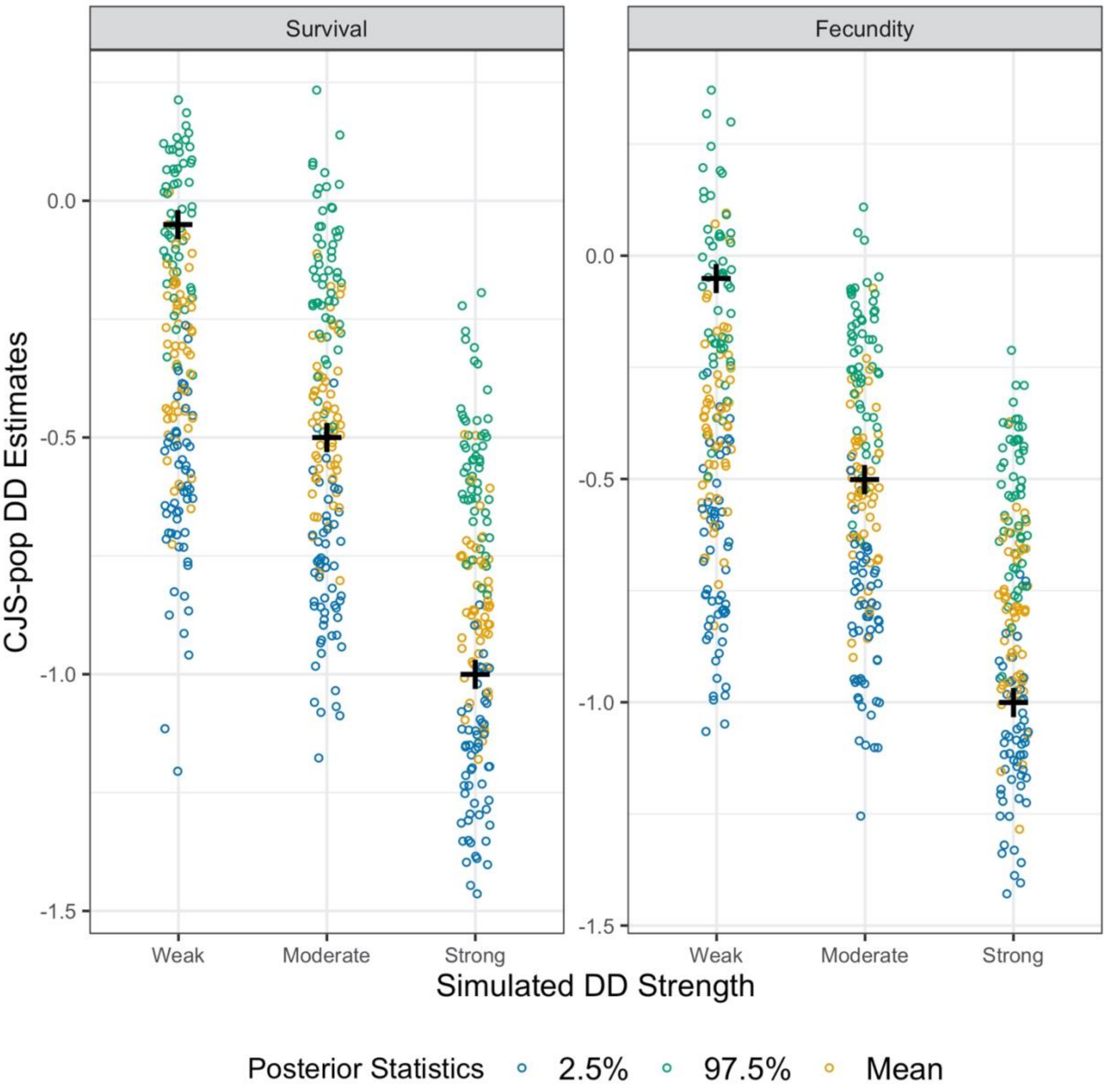

**Figure S1:** Posterior statistics of Density Dependence in Survival and Fecundity in simulated data sets with 10 populations and carrying capacity of 150. “+” sign shows the true value of the density dependence parameter used to generate the simulation set. If the “+” sign is centered on orange dots (mean of the posterior distribution) density dependence parameters in that simulation set is estimated without bias. If the “+” sign is placed around blue dots (2.5% quantile of the posterior distribution) density dependence is underestimated, or conversely if it is placed around green dots (97.5% quantile of the posterior distribution) density dependence is overestimated.

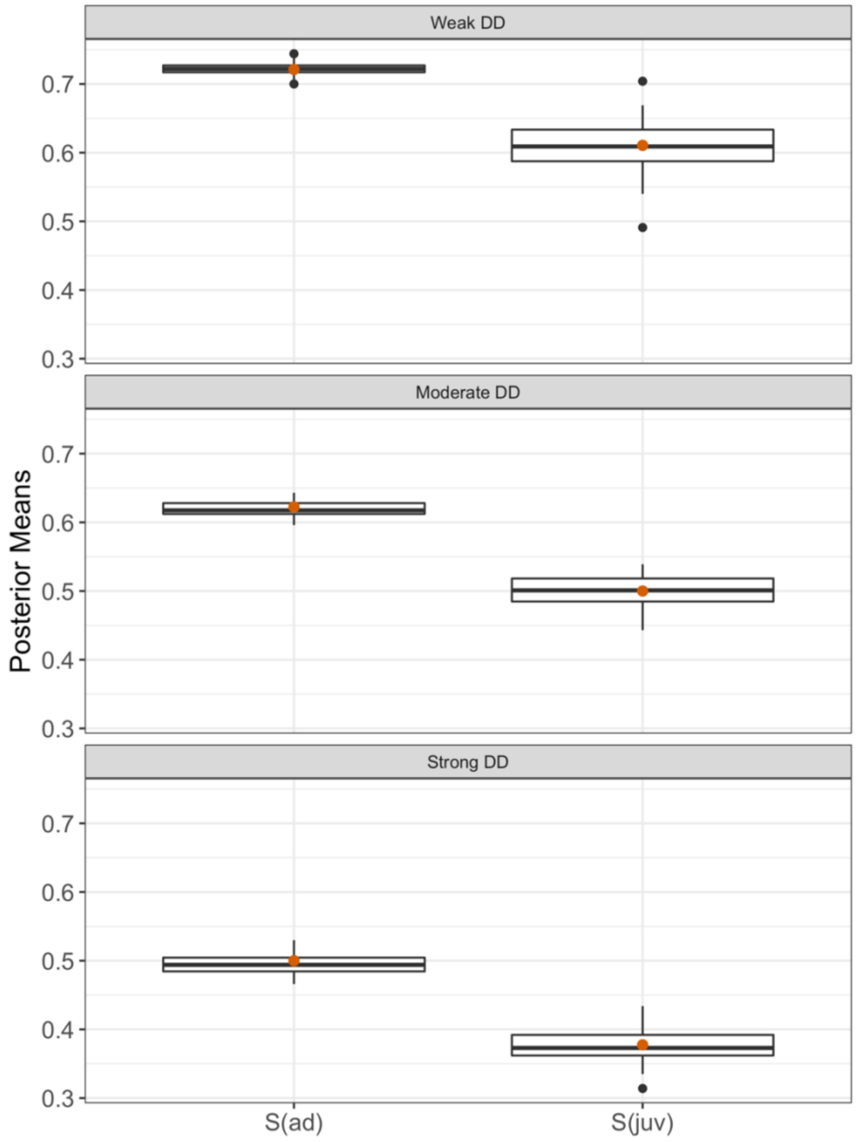


**Figure S2:** Boxplots of posterior means of survival parameters estimated by CJS-pop fit to simulated data with 10 populations and a carrying capacity of 150. Orange dots are true parameter values used in simulations to generate data. S(ad) and S(juv) are adult and juvenile survival rates at mean population size, respectively.

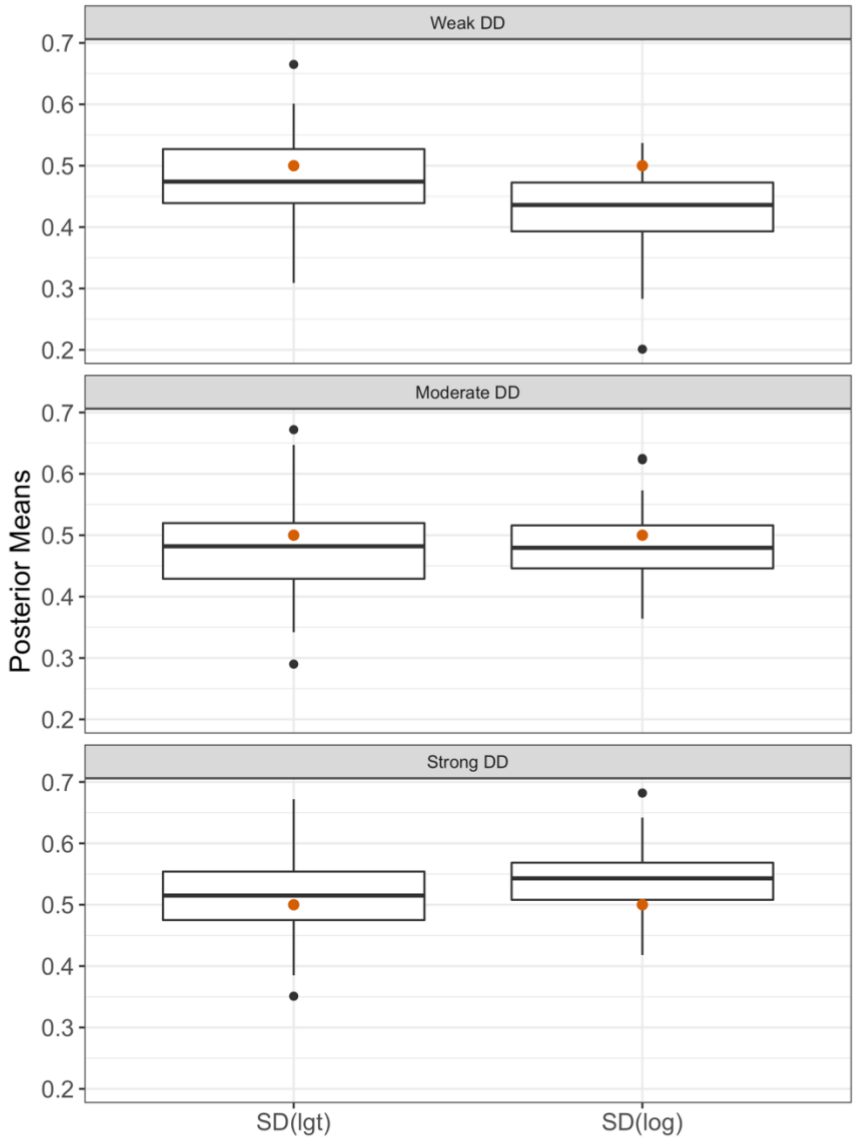


**Figure S3:** Boxplots of posterior means of process variance estimated by CJS-pop fit to simulated data with 10 populations and a carrying capacity of 150. Orange dots are true parameter values used in simulations to generate data. SD(lgt) is the square root of process variance of survival in logit scale and SD(log) is the square root of process variance of fecundity in log scale.

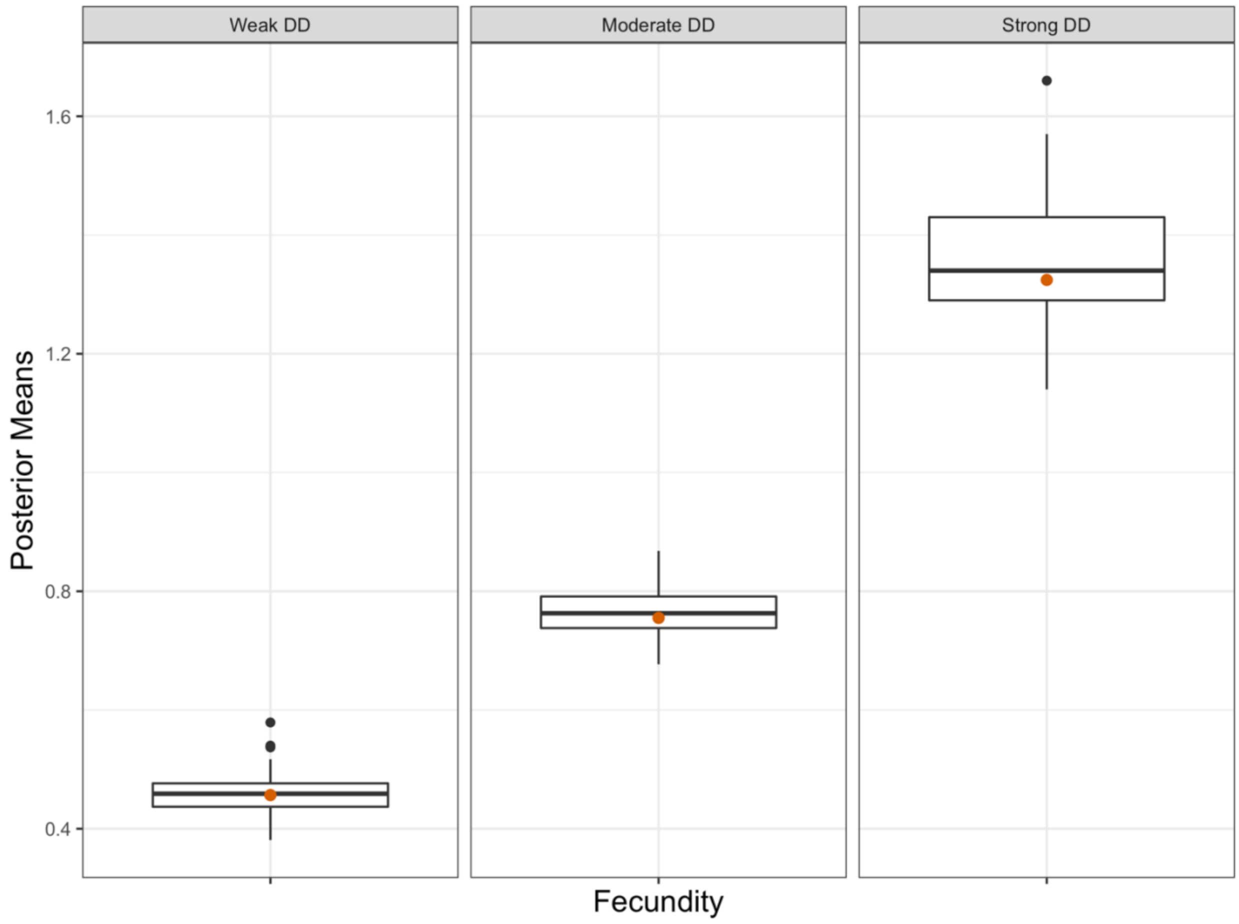


**Figure S4:** Boxplots of posterior means of fecundity at mean population size estimated by CJS-pop fit to simulated data with 10 populations and a carrying capacity of 150. Orange dots are true parameter values used in simulations to generate data.

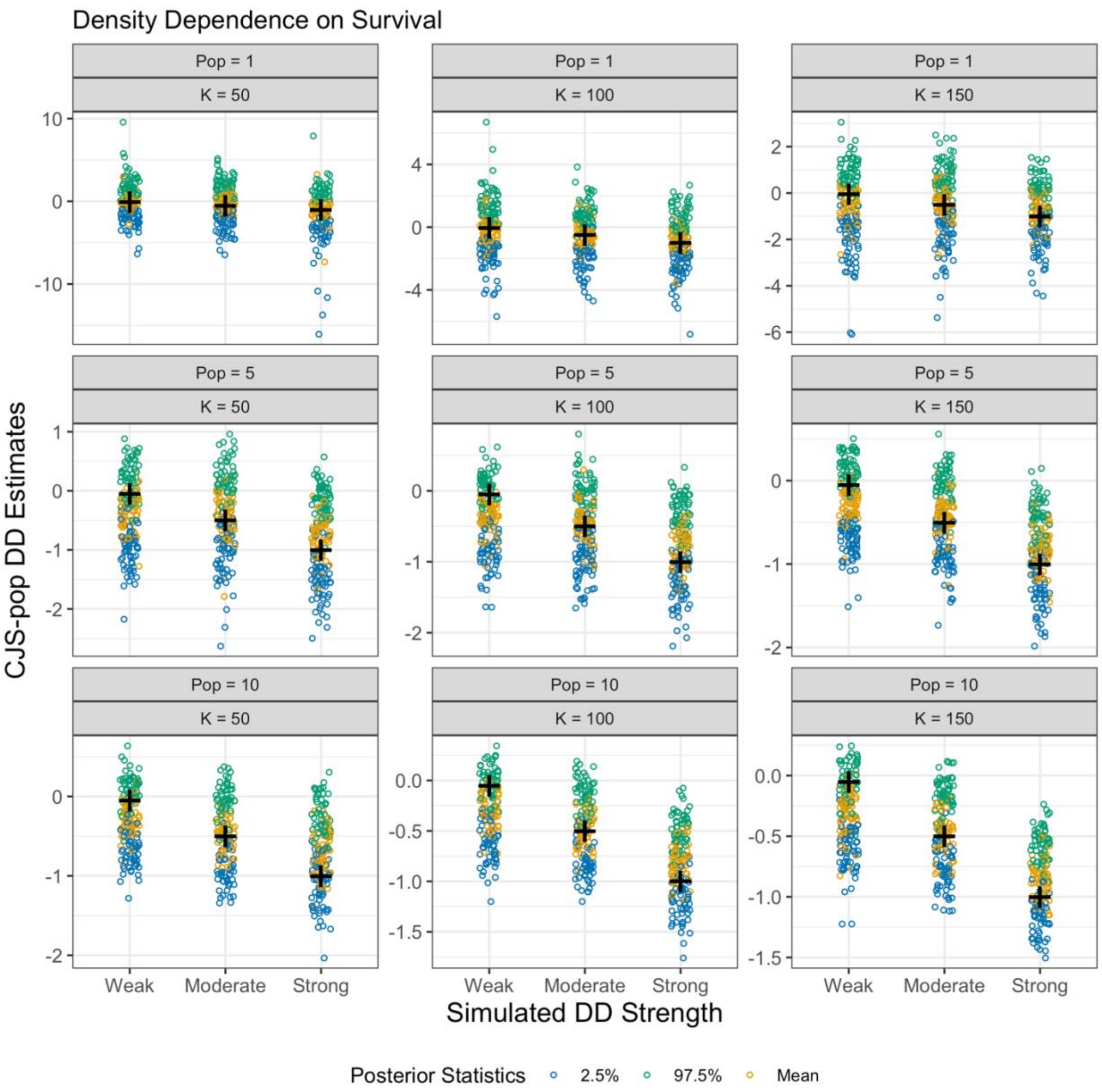


**Figure S5:** Posterior statistics of Density Dependence on Survival of each simulation set. "+" sign shows the true value of the density dependence parameter used to generate a simulation set. If the "+" sign is centered on orange dots (mean of the posterior distribution) density dependence parameters in that simulation set is estimated without bias. If the "+" sign is placed around blue dots (2.5% quantile of the posterior distribution) density dependence is underestimated, or conversely if it is placed around green dots (97.5% quantile of the posterior distribution) density dependence is overestimated.

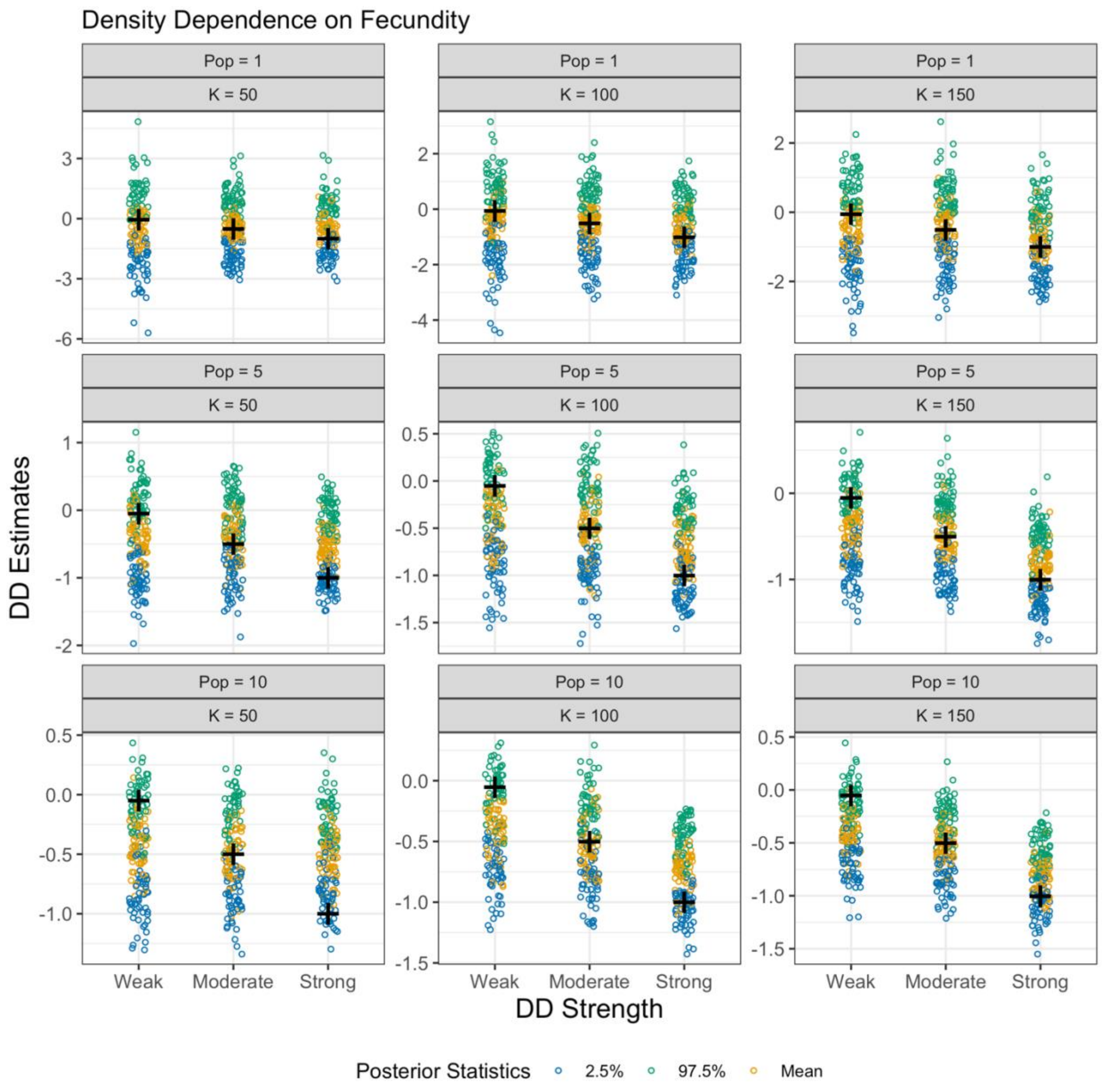


**Figure S6**: Posterior statistics of Density Dependence on Fecundity of each simulation set. "+" sign shows the true value of the density dependence parameter used to generate a simulation set. If the "+" sign is centered on blue dots (mean of the posterior distribution) density dependence parameters in that simulation set is estimated without bias. If the "+" sign is placed around red dots (2.5% quantile of the posterior distribution) density dependence is underestimated, or conversely if it is placed around green dots (97.5% quantile of the posterior distribution) density dependence is overestimated.

**Table S1:** Posterior statistics of parameters estimated with density dependent CJS-pop fit to Brown Creeper data. Rhat is the multivariate version of Gelman and Rubin's convergence diagnostic, and n.eff is the effective sample size of posterior distributions (out of 10000). S (ad) and S (juv) are adult and juvenile survival rates at carrying capacity, respectively; $\beta$ is density dependence strength on survival in logit scale; $\sigma_s$ is the process variance of survival in logit scale; F is fecundity at carrying capacity; $\zeta$ is density dependence on fecundity in log scale; $\sigma_f$ is the process variance of fecundity in log scale; P (ad) and P (juv) are monthly capture probabilities of adults and juveniles, respectively; $\pi$ (ad) and $\pi$ (juv) are the probabilities of adult and juvenile individuals being residents, respectively; $\rho$ (ad) and $\rho$ (juv) are the probabilities correctly identifying true adult and juvenile residents as residents, respectively.

| | MEAN | 2.5% | 97.5% | RHAT | N.EFF |
|---|---|---|---|---|---|
| **S (AD)** | 0.422 | 0.347 | 0.500 | 1.00 | 2716 |
| **S (JUV)** | 0.315 | 0.19 | 0.482 | 1.00 | 921 |
| $\boldsymbol{\beta}$ | -0.273 | -0.735 | 0.219 | 1.00 | 5520 |
| $\boldsymbol{\sigma_s}$ | 0.228 | 0.010 | 0.608 | 1.00 | 1881 |
| **F** | 1.940 | 1.210 | 3.010 | 1.01 | 911 |
| $\boldsymbol{\zeta}$ | -0.136 | -0.407 | 0.133 | 1.00 | 7951 |
| $\boldsymbol{\sigma_f}$ | 0.974 | 0.843 | 1.120 | 1.00 | 6355 |
| **P (AD)** | 0.047 | 0.039 | 0.055 | 1.00 | 3889 |
| **P (JUV)** | 0.012 | 0.008 | 0.018 | 1.00 | 992 |
| $\boldsymbol{\delta}$ | 0.660 | 0.440 | 0.885 | 1.00 | 6075 |
| $\boldsymbol{\pi}$ **(AD)** | 0.630 | 0.465 | 0.850 | 1.01 | 1258 |
| $\boldsymbol{\pi}$ **(JUV)** | 0.248 | 0.145 | 0.409 | 1.00 | 1021 |
| $\boldsymbol{\rho}$ **(AD)** | 0.144 | 0.100 | 0.198 | 1.00 | 1744 |
| $\boldsymbol{\rho}$ **(JUV)** | 0.085 | 0.043 | 0.147 | 1.00 | 1674 |
