## Appendix S2 for "Estimating fecundity and density dependence from mark-recapture data for making population projections"

### Appendix S2: Model and Simulation Details

#### Contents

1. Details of the residency model
2. Variance structure of CJS-pop
3. Double estimation of the expected population size of juveniles
4. Simulating Capture Histories
5. Software
6. Priors
7. Goodness-of-fit testing
8. Model Assumptions
9. Comparison to previous model results
10. Population sizes as expected values vs random variables
11. Supplementary References

**1. Details of the residency model**

We employ a simplified version of the models described in Saracco et al. (2010) when modelling residents. This approach accounts for transients that will leave the population after their first capture but does not model movement between populations. We assume that there are two parameters, $\rho_1$ for juveniles and $\rho_2$ for adults, that governs the probability that we can correctly categorize a true resident as a pre-determined resident. The residency data (1 for pre-determined residents, 0 for potential transients), $r_i$, is a Bernoulli random variable and is conditional on the $i$th individual being a true resident ($R_i$):

$$r_i \sim \text{Bernoulli}\left(R_i \cdot \rho_{S_{i,f_i}}\right) \quad (1)$$

where, $f_i$ is the first capture year of the $i$th individual, and $S_{i,f_i}$ corresponds to the stage of the $i$th individual in its first capture year. We also assume that there is a parameter for each stage, $\pi_1$ and $\pi_2$, that governs the probability that a juvenile or an adult in the population is a true resident, respectively. $R_i$ is a Bernoulli random variable with probability $\pi_{S_{i,f_i}}$:

$$R_i \sim \text{Bernoulli}\left(\pi_{S_{i,f_i}}\right) \quad (2)$$

$$Z_{i,(t+1)} \sim \text{Bernoulli}\left(Z_{i,t} \cdot \phi_{(S_{i,t}),(g_i),t} \cdot R_i\right) \quad (3)$$

This model effectively filters out some of the capture histories that only has the first capture followed by no captures for the remainder of study period (a single 1 and trailing 0s in their capture history). If true residency of an individual ($R$) is estimated to be 0 (a transient), then its state ($Z$), will also be automatically 0 (see eq. 5) and it will not lower the survival estimate. $\pi$ and $\rho$ are nuisance parameters and are not used in projections explained below.

r is the data related to residency. We denote an individual pre-determined resident (1 in the data) if it was re-captured at least 10 days after its first capture in its first year. The assumption here is that if it didn't leave the population in its first year and was recaptured, then it must be a resident. It's called pre-determined resident because this classification is done before building the model. If it did leave the population and was not captured again in its first year of capture, then there is a chance that it could be a transient and may have simply left the population, or it could be a resident and will be recaptured in the following years. These are denoted as potential transients (0 in the data).

R is a latent residency variable, the “true” state of an individual in terms of its residency. Due to the model structure if r is 1 then R is automatically 1 as well; and if an individual's R is 0, that individual cannot be mistakenly categorized as a pre-determined resident. So, this model re-classifies potential transients as either true residents or true transients. With this model, if R is 1 for an individual (but we don’t know this since this is a latent state) then there is nevertheless a chance that that individual will be wrongly categorized as a potential transient for the simple fact that it was not recaptured again in its first year. This is the probability $\rho$. Estimation of this probability makes it possible to estimate $\pi$, the probability that an individual in a population is a resident (as opposed to a transient). Estimation of these two probabilities by using r allows estimation of the latent state R, which is used in survival models to filter out some individuals estimated to be transients. This partially accounts for apparent survival estimates. The product $\rho \cdot \pi$ is equal to the number of pre-determined residents divided by the total number of individuals (this statement is true for juveniles and adults, separately). This approach is similar to how survival rates, capture probabilities, and Z (the latent state of an individual being alive and, in the population,) are modeled together.

**2. Variance structure of CJS-pop**

We used temporal random effects when modelling survival for Brown Creeper. This approach assumes that populations are spatially fully correlated in their temporal variability (i.e., all populations experience “good” years or “bad” years at the same time); this assumption becomes especially important when survival is also modeled as a function of population-level covariates (here, a density index). We tried modelling spatial autocorrelation in temporal response for both survival and fecundity so that closer populations in space would have more similar temporal random effects. We could not get this model to converge, however. We also tried the opposite approach, where there is no spatial autocorrelation in temporal random effects (i.e., each population in each year can have different and independent random effects). We call this spatio-temporal variability. This did not work for survival; we think at least for Brown Creeper, there is not enough information in the capture history for spatial variability of survival. However, this improved the fecundity model, which becomes problematic with temporal-only random effects, in terms of convergence issues and low effective sample size of MCMC chains. So, survival is modeled with temporal random effects, and fecundity with spatio-temporal random effects.

We can show that the estimated value of $\sigma_f^2$ for Brown creeper is high by generating fecundity values at mean population size by using the $\theta$ estimate (Table S1; average fecundity across populations and years) and add 10000 randomly sampled effects from $N(0, \sigma_f^2)$. The 95% credible interval of the exponent of these generated values (because they were in log scale) was 0.29 and 13.09. Because we don't limit this distribution, in very rare cases values as high as 55 can be sampled, which would not be plausible for this species.

In applications of  CJS-pop to specific cases, this uncertainty in fecundity estimates can be reduced by removing outliers. For example, Ryu et al. 2016 proposed a similar framework, in which they removed populations that had fewer than 5 adult captures. Outliers can also be removed by running CJS models first and removing populations with very low capture probability, which may cause unrealistically high juvenile population size estimates. Another option is to calculate fecundity without running any model, by dividing the captured number of juveniles with adults, and removing outlier fecundity estimates. These outlier estimates will remain as outliers even after fitting CJS-pop because capture probability model of CJS-pop does not specifically account for populations and years that has uncharacteristically high juvenile captures compared to adults. These populations and years with high number of juvenile captures will lead to even higher fecundity estimates if on average juvenile capture probability is lower than adult capture probability. High fecundity estimates in some populations and years also results in high process variance estimates ($\sigma_f^2$). Removing these estimates from the analysis would reduce both $\theta$ and $\sigma_f^2$. It is also possible to add a temporal random effect term to the capture probability model, separately for adults and juveniles, in order to account for years that seemingly has higher juvenile capture probability compared to adults. We preferred not to use any of these approaches for this manuscript because we aim to present a more fundamental form of CJS-pop.

In this implementation of CJS-pop, we did not include spatial or temporal auto-correlation in vital rates. Our attempts at doing so led to convergence problems. We also assumed that random temporal effects on survival and on fecundity were independently distributed. Although covariances between vital rates can be explicitly modeled in CJS-pop (Elderd and Miller 2015), this would likely cause convergence problems as well.

**3. Double estimation of the expected population size of juveniles**

Cyclical relationships usually point to model specification issues. Here, we know which part of the model causes a cyclical relationship. When calculating density index of each year, we standardize that year's population size with average size of the population across 17 years (or less if sampling interval was shorter). $n_{1,k,t}$ and the numerator in equation 3 in the main text is compatible when modelling fecundity, because we use density index $D_{k,(t-1)}$ so the time index in the numerator is also t-1. However, denominator in equation 3 (the population average over T years) has information coming from year t because $n_{1,k,t}$ is used to calculate expected juvenile population size in equation 2 in the main text. From the perspective of JAGS, $n_{1,k,t}$ is used to estimate $D_{k,(t-1)}$, which is in turn used to estimate $N'_{1,k,t}$. Finally, $n_{1,k,t}$ is modelled as a Poisson random variable whose expected values is determined by $N'_{1,k,t}$, hence the cyclical relationship. We don't think this cyclical relationship is a model specification issue. The long-term population average is used in place of a carrying capacity and any one $n_{1,k,t}$ is only a small part of a $D_{k,(t-1)}$ estimate.

#### 4. Simulating Capture Histories

##### 4.1 The Simulation Framework

We simulated several sets of capture histories in order to test the CJS-pop's ability to correctly retrieve true parameter values, and in order to uncover any inherent biases especially when quantifying density dependence (DD) strength.

Fundamentally, the simulations are typical stage-structured models with two stages (adults and juveniles):

$$\begin{bmatrix} N_{1,k,(t+1)} \\ N_{2,k,(t+1)} \end{bmatrix} = \begin{bmatrix} \phi_{1,t} * F_{k,(t+1)} & \phi_{2t} * F_{k,(t+1)} \\ \phi_{1,t} & \phi_{2,t} \end{bmatrix} \cdot \begin{bmatrix} N_{1,k,t} \\ N_{2,k,t} \end{bmatrix} \quad (4)$$

where $N_{1kt}$ $N_{2kt}$ are juvenile and adult populations sizes of population $k$ and time $t$, respectively. Survival and fecundity are modeled as functions of density:

$$logit(\phi_{1,t}) = \alpha_1 + \beta \cdot D_{k,t} + \epsilon_{k,t} \quad (5)$$

$$logit(\phi_{2,t}) = \alpha_2 + \beta \cdot D_{k,t} + \epsilon_{k,t} \quad (6)$$

$$log(F_{k,(t+1)}) = \theta + \zeta \cdot D_{k,t} + \omega_{k,t} \quad (7)$$

$$\epsilon_{k,t} \sim Normal(0, \sigma_s^2) \quad (8)$$

$$\omega_{k,t} \sim Normal\left(0, \sigma_f^2\right) \quad (9)$$

The parameters $\alpha_1$, $\alpha_2$, and $F_{k,(t+1)}$ have different interpretations than their respective models in CJS-pop because we are using density directly as a covariate (as opposed to subtracting 1 in CJS-pop). These parameters now represent theoretical maximum survival and fecundity values of a species when a population is at 0 density. Density itself is calculated by dividing population size at each time step to a set carrying capacity. We assumed spatio-temporal variance for both fecundity and survival.

Additionally, however, we followed the fate of each individual as they entered a population. This was necessary to create a unique capture history of an individual if it was ever captured. Each simulation consisted of multiple closed populations so there was no emigration or immigration between them. Multiple populations served as a way to control sample size but also to make data sets more similar to MAPS capture histories which consist of multiple locations. We assumed constant carrying capacity across years. We also assumed constant effort and therefore a constant capture probability across years for adults and juveniles. In all of our simulations we set monthly adult capture probability to 0.1 and juvenile capture probability to 0.02.

We set up a simulation scheme where we explored the effect of sample size on quantifying density dependence strength. Sample size in a CJS-pop is determined by three factors: 1) Time series length, 2) Number of populations, and 3) Size of the populations. Here we set the time series length to 17 years, which is the maximum time series length for Brown Creeper in MAPS data set we're using. We simulated three cases with 1, 5, and 10 populations, and three carrying capacities (which controls population size): 50, 100, 150. For each combination of the number of populations and carrying capacity, we generated capture history data sets using weak, moderate and strong density dependence strength on survival and fecundity, which created 27 separate simulation sets (for example 1 population with carrying capacity of 100 and low density dependence, or 10 populations, each with carrying capacity of 150 and high density dependence). For each simulation set, we generated 56 capture history data sets and fitted CJS-pop to each one. The number of capture history data sets was a trade-off between computational time and sample size.

Density dependence strength refers to the value of $\beta$ and $\zeta$ in CJS-pop framework. We set the weak strength to be $-0.05$, moderate strength to be $-0.5$, and strong strength to be $-1$. We determined these by calculating the growth rate ($\lambda$) of the population at 0 density, which is the maximum growth rate of the population ($R_{max}$=exp($r_{max}$)). $R_{max}$ can be calculated for a set of vital rates by employing eq. 1 (but by not setting it to 1). Accordingly, $R_{max}$ for weak, moderate, and strong density dependence were 1.03, 1.5, and 2.97, respectively. Here, we used the allometric relationships reviewed and reported in Figure 2c of Fagan et al. 2010. They report a metric equivalent to $r_{max}$ across different taxa including birds based on count data. Among birds with the size of Brown Creeper (~100 g), they report no species with $r_{max}$ as low as 0.02 (i.e., Rmax=1.03). However, allometric relationships are variable, and it is biologically possible to have even stronger DD for species that is the size of songbird. While we limited our simulations to three values of DD strength, additional simulations for detecting bias patterns are necessary, especially if CJS-pop is going to be applied for different taxa.

Across all CJS-pops that were fit to different combinations of population, carrying capacity and density dependence strength, we removed from the analysis any model (i) with at least one parameter that had R-hat value larger than 1.05, or (ii) with at least one p-value $> 0.95$ or $<$ 0.05.

##### 4.2 Extending the simulation framework

We believe simulations with more parameter combinations is a future research topic for CJS-pop. For example, we did not explore how CJS-pop would perform with a single population that has a larger carrying capacity than 150.  A larger number of individuals in a population might give better results than what is reported for single population simulations in the main text, but we believe this will still lead to high uncertainty for density dependence (DD) estimates. This is because a single population simulation with environmental stochasticity gives only a single population trend in which the effects of density dependence might be dominated by stochasticity. Multiple populations, if temporal variation in their vital rates is at least partially independent, provide different population trajectories which in turn should allow CJS-pop to estimate DD with less uncertainty and bias. Single population data or multiple population data with high spatial autocorrelation can potentially estimate DD with lesser uncertainty than what is reported in the main text, if the models include environmental covariates that affect vital rates. This is because these covariates should account for some of the stochasticity in the trajectory, therefore allowing the model to pick up a signal for DD. A promising future direction is developing a simulation framework that incorporates spatial-autocorrelation and environmental factors and explores how their inclusion and omission affects CJS-pop estimates.

#### 5. Software

All data generation, manipulation, and analysis were done with R 3.5 (R Core Team, 2018). We with R packages R2Jags (Su and Yajima, 2015) and rjags (Plummer, 2016). Post-hoc data wrangling was done using the R package MCMCvis (Youngflesh 2018). Rhat and effective sample size of MCMC chains is calculated by MCMCvis package, which follows Brooks and Gelman (1998).

**6. Priors**

We used weakly informative priors for all parameters in the framework. Gelman (2006) defines a prior distribution as weakly informative if it is proper and the information it provides is intentionally weaker than actual available prior knowledge. In the same vein, in terms of the information it encodes, a weakly informative prior is considered to be between objective and expert priors (Simpson et al. 2017), or structural and regularizing priors (Gelman et al. 2017).

Additionally, we used the following growth rate ($\lambda$) equation to set the priors of vital rates:

$$S_A + S_J * F = 1 \tag{10}$$

where, $S_A$ and $S_J$ are average survival rates of adults and juveniles at carrying capacity and $F$ is the fecundity at carrying capacity. This equation states that population growth rate ($\lambda$) is equal to 1 at carrying capacity. The degrees of freedom for these three parameters is 2. If we know the estimates of two of these parameters, the third one can be directly calculated. Among these three, juvenile survival is the hardest to estimate because juvenile recaptures are rare. So, we only used priors for adult survival and fecundity and calculated juvenile survival as:

$$S_J = \frac{1 - S_A}{F} \tag{11}$$

In the density independent models in which we modeled only the average vital rates we assumed all three rates were independent and did not use this relationship.

The list of all priors used for CJS-pop fit to Brown Creeper data is as follows:

$$\begin{aligned}
S_{ad} &\sim U(0,1)\\
\beta &\sim t(0,10,1)\\
\sigma_s &\sim t(0,10,1)[0,\infty]\\
\gamma_{ad} &\sim t(0,10,1)\\
\gamma_{juv} &\sim t(0,10,1)\\
\delta &\sim t(0,2.5,1)\\
F &\sim N(0,10)[0,\infty]\\
\zeta &\sim t(0,5,1)\\
\sigma_f &\sim t(0,5,1)[0,\infty]\\
\pi_{ad} &\sim U(0,1)\\
\pi_{juv} &\sim U(0,1)\\
\rho_{ad} &\sim U(0,1)\\
\rho_{juv} &\sim U(0,1)
\end{aligned}$$

*t* states a t distribution where the first argument is the mean, the second is the standard deviation, and third is degrees of freedom. *N* is normal distribution where the first argument is the mean, and the second is standard deviation. *U* is the uniform distribution where arguments indicate the low and high limits of the distribution. "[,]" is used to indicate between which limits a distribution was truncated.

**7. Goodness-of-fit testing**

Our model has more assumptions than a typical CJS model (see below), which, if violated, might lead to a bad model fit. We test the goodness-of-fit (GoF) of this framework with posterior predictive checks via Bayesian p-values (Kéry and Royle 2016). In a posterior predictive check, new data are generated conditional on the estimated parameters in each iteration of the MCMC, and these new data are compared to the original data used to fit the model. A similar logic of the frequentist p-value applies here as well, the probability of getting as or more extreme data conditional on the model. The goal is to calculate $P(T(y^{new},\theta) \geq T(y,\theta))$, where $T(y,\theta)$ is a summary statistic calculated both for new data and the observed data, and $\theta$ here is used to represent the vector of parameters.

We chose Freeman-Tukey (FT) statistic as the test statistic for both the capture history component and fecundity (Conn et al. 2018). FT can be calculated as $\sum\left(\sqrt{y_i} - \sqrt{E(y_i)}\right)^2$, where $E(y_i)$ is the expected value. We pool all the secondary periods within their primary period when calculating the FT statistic. The expected value then becomes the number of expected recaptures of an individual within a year, which can be calculated as $\sum_{h=1}^{H} p_{ith}$. For fecundity, the expected value is the expected number of captured juveniles, which is equal to $P_{2kt} \cdot N'_{2kt}$ for year $t$ and population $k$.

The interpretation of a Bayesian p-value is different from a frequentist p-value. Values closer to 0.5 indicate a better fit, and values too close to 0 or 1 show a lack of fit. There are no widely accepted thresholds, however (Kéry and Royle 2016). We set 0.05 and 0.95 as lower and higher thresholds, respectively, and state that any model outside of these thresholds do not represent the data well.

**8. Model Assumptions**

The survival and capture probability part of this framework is a Cormak-Jolly-Seber model (CJS) with the addition of density and effort as covariates. See Lebreton et al. (1992), Williams et al. (2002), and Cooch and White (2016) for a more in depth discussion of CJS model assumptions.

However, this analysis also employs a robust design, so it is accompanied by an additional set of assumptions, especially for the sub capture occasions (secondary period). Within each primary capture period, population is assumed to be closed, there is no emigration and no deaths. Because we set this period to be the whole breeding season (4 months) it is very likely that this assumption is violated in this framework. Some individuals may die or leave the population in a given secondary period and this would lead to a biased estimate of capture probability. This is important because capture probability is not a nuisance parameter in this framework and we use it explicitly to estimate true population size of adults and juveniles in a given year and population. Immigration and births also violate the closure assumption, but this is not of vital importance because estimation of survival rates and capture probabilities are conditional on first capture.

In our opinion, this type of violation of the population closure assumption is acceptable in this framework. Capture probability estimates, if biased due to the reason explained above, are lower than their true value. This leads to an inflated population size estimate under the heuristic population size estimator, but not necessarily to an inflated density estimate. This is because density is calculated by standardizing the population size estimate of each year with the mean of population size across all sampling years. Any bias in population size in a given year would also be prevalent in the mean population size. The only deviation from this expectation would be when capture probability of juveniles and adults are largely different, and when there are considerably more juveniles than adults in a population. In that case the estimate of density dependence strength might be biased, a lower estimate compared to its true value. However, this also means that if we find a signal for density dependence it is likely not a false positive, whereas a lack of signal might indicate a false negative due to the structure of the data and the framework.

Biases in capture probability has a potential to affect fecundity estimates more so than strength of density dependence. This is because not all juveniles are available for capture across a whole primary period. If they leave the population within a primary period the capture probability estimate would be deflated, and this is also true for adults; but uniquely for juveniles they may not be available for capture simply because they were not born early in the breeding season. Probability of capturing a juvenile that was born in May at least once is higher than a juvenile that was born in July. However, using the heuristic population size estimator means that we assume all were born in May. So even if capture probability for a secondary period is perfectly estimated the estimate of true population size of juveniles can still be deflated if there are many juveniles born late in the breeding season.

This means that there are two bias factors operating on the opposite direction in estimating true juvenile population size and therefore fecundity. When Juvenile leaves within a primary period, this results in a lower estimate of capture probability leading to a higher juvenile population size and fecundity. If many juveniles are born late in the breeding season, the capture probability estimate is not affected but due to the assumptions of the heuristic population size estimator, juvenile population size and fecundity estimates are biased and are lower than the true value. It is impossible to exactly discern which of the two factors are more prevalent in any given data set.

A similar situation is caused by adults that enter a population by immigration. Depending on when they immigrate adults may not be available for capture across a whole primary period. If there are many late immigrants, this would also deflate the adult population size estimate. Again, two opposing forces affecting the adult population size: early emigrants would deflate capture probability leading to an inflated adult population size estimate, while late adult emigrants lead to a deflated estimate.

As argued above, whatever the bias factor is, it is prevalent both in the yearly population size estimate and the mean population size estimate. So, we expect their effect on the estimation of density dependence to be minimal. However, strong temporal variation in the severity of this bias would affect density dependence estimates. While we don't suspect this is the case for Brown creeper, this variation can be caused by ecological reasons, due to the variation in survival or fecundity, or it can be caused by the methodology, as in there is strong year to year variation in the capture probability of individuals. Additionally, it is also important to mention that the heuristic population size estimator has a large variance. A large error in estimating population size might lead to a reversal of yearly trends. A slight decrease in population size from year $t$ to $t+1$ might be estimated as an increase. This type of trend reversal makes it especially hard to correctly estimate the density dependence strength. If these reversals are excessive and without bias, the estimated strength should simply be centered on or close to 0.

**9. Comparison to previous model results**

CJS-pop's vital rate estimates of Brown Creeper are similar to those of a previous CJS model built with the same data that also accounts for transients (DeSante et al. 2015). This previous study found adult survival to be 0.389, comparable to, but slightly lower than, CJS-pop's estimate of 0.422 (Table S1). The difference is likely due to the different structure of our model, which also estimates the dependence of survival on population density. Capture probabilities estimated in CJS-pop were lower compared to DeSante et al. (2015). We estimated 0.047 and 0.012 monthly capture probability for adults and juveniles, respectively (Table S1). This corresponds to 0.17 yearly capture probability for adults, which was found to be 0.23 in their CJS model.

#### 10. Population sizes as expected values vs random variables

In a typical IPM, population size is estimated as a binomial and a Poisson process. For example, from Ahrestani et al. 2017 with slight modifications:

$$S_{s,(t+1)} \sim Binomial(\phi_{s,t}, N_{s,t}) \quad (12)$$

$$G_{s,(t+1)} \sim Poisson(\gamma_t N_{s,t}) \quad (13)$$

$$N_{s,(t+1)} = S_{s,(t+1)} + G_{s,(t+1)} \quad (14)$$

where $N_{s,t}$ is the population size in time t and population s; $\phi_{s,t}$ is the annual survival probability from year t to t+1; $S_{s,(t+1)}$ is the number of individuals survived from year t to t+1; $\gamma_t$ is the recruitment to a population from year t to t+1; and $G_{s,(t+1)}$ is the number of recruited individuals from year t to t+1. In an IPM, $N_{s,t}$ would also be modeled separately using population count data. Using information from population count data and mark-recapture data at the same time, an IPM would ensure that vital rate estimates are compatible to both data sets. We tried a similar approach, but our models did not converge. This is because population count data helps with the convergence of the whole framework in a complex model such as ours.

We see using the expected value vs. population count data to model and estimate population sizes as trade-offs. Modelling population processes explicitly is preferable as it is a better representation of a natural process in statistical form. However, this approach requires additional data in the form of population counts and it cannot be applied when data are the limiting factor. Estimating population size as expected values ignores the uncertainty arising from population processes and it employs only a point estimate. However, point estimates can still be used when the only available data are robust design mark-recapture data, so this can be useful when data are limited.
