## Appendix S3-S4 for "Estimating fecundity and density dependence from mark-recapture data for making population projections": AppendixS3.Rmd

````
---
title: "CJS-pop models for MAPS Data"
output:
  html_document:
    df_print: paged
---

### Introduction

CJS-pop models for analysing capture history data from the MAPS program. See this **Sen and Akcakaya (in review)** for detailed model description.

JAGS language is not very versatile and seemingly straightforward tasks in other languages may require a few extre lines in JAGS. Here we explain some of the programming tricks we used in the models below.

1. **Fastening the loops**. For each population, models run only from the first year to the last year sampling took place in a population. So, while the whole study as a whole is between 1992 to 2008, that is not the case for most populations. Survival and capture history models are conditioned on the first capture of an individual but end at the last year of the individual's population, not necessarily extending to 2008. Population size and density are only estimated between first and last year of captures of each population. This way loops do not extend unnecessarily into early or late years with no capture effort in populations.

2. **Imputing missing data (Part 3.2)**. As explained above density is not estimated for before the first year of capture or after the last year of capture in a population. However, between first and last years there might still be years with no capture effort, and hence 0 capture probability (or rather extremely low capture probability) and 0 captured individuals, making the estimation of expected population size difficult. We impute these years with a density value of 1 (in other words a density index of 0 after standardization). We are assuming that the population is stable and is at carrying capacity in years with no available infornation. 

On a similar note, rue to the asymptotical structure of logistic regression we never estimate 0 capture probability even when there is no effort, but rather a number too low. This can be an issue when calculating average population size because of the correction we do when estimating the expected population size, and can lead to abnormally large population size estimates when there is no capture effort. To circumvent this issue we impute years with no capture effort with a population size of 0. This does not bias density estimates because only years with capture effort are considered when estimating density, as explained below.     

3. **Density Estimation (Part 3.2)**. When estimating density we only consider years when there is capture effort. We count these years in a matrix called "n1", whose rows are populations and columns are years. We use the equal function to see if the total capture effort is equal to 0 (no capture effort), and if it is, equal function assings 1 to the corresponding matrix element, or 0 if it is not. Summing across a single row of the n1 matrix (between first and last capture years) counts the number of years with no effort. We substract this number from the range of years (last year - first year + 1) in that population to count the total number of years with capture effort in a population. This information is assigned to the vector "n2". Total population size is divided by n2 to calcualte average population size across years, which in turn used to estimate density in each year in each population.

4. **Indexing Fecundity Data (Part 4)**. We filter fecundity data so that only years and populations that have at least 1 adult are included in the final data. This means that some years and populations will be missing from this data, so fecundity model can not be covered with a regular "for loop" across years and populations. Instead we use two vectors "pop_index"" and "year_index" that specifies which element in the fecundity data, itself also a vector, belongs to which population and year. This way a single "for loop" can loop through the fecundity data, pop_index, and year_index while still subsetting correct elements in the density matrix with pop_index and year_index.

The first model below corresponds to the model presented in **Sen and Akcakaya(in review)**. Rest of the models are similar in structure with different variations and sometimes with different tricks. We do not explain those in detail. 

### 1. Density dependent CJS-pop with residents

```{r, message=FALSE}

{
sink("cjspop.jags")
  cat("
      model {
      
      # Part 1: Priors
      
        # Part 1.1: Priors for survival model

        survival_ad ~ dunif(0,1)
        survival_juv <- (1-survival_ad)/fecundity
        alpha[2] <- logit(survival_ad)
        alpha[1] <- logit(survival_juv)
        beta ~ dt(0,pow(10,-2),1)
        sigma_s ~ dt(0,pow(10,-2),1)T(0,)
        tau_s <- 1/(sigma_s^2)

        for (t in 1:(nyear-1)) {
          epsilon[t] ~ dnorm(0, tau_s)
        }

        # Part 1.2: Priors for capture porbablity model

        gamma[1] ~ dt(0,pow(10,-2),1)
        gamma[2] ~ dt(0,pow(10,-2),1)
        delta ~ dt(0,pow(2.5,-2),1)
      
        # Part 1.3: Priors for fecundity model

        fecundity ~ dnorm(0,pow(10,-2))T(0,)
        theta <- log(fecundity)
        zeta ~ dt(0,pow(5,-2),1)
        sigma_f ~ dt(0,pow(5,-2),1)T(0,)
        tau_f <- 1/(sigma_f^2)
        psi ~ dunif(0,1)

        for (l in 1:nfec) {
          omega[l] ~ dnorm(0, tau_f)
        }
        
        # Part 1.4: Priors for residency model
      
        pi[1] ~ dunif(0,1)
        pi[2] ~ dunif(0,1)
        rho[1] ~ dunif(0,1)
        rho[2] ~ dunif(0,1)

      # Part 2: Robust design mark-recapture model

        # Part 2.1: Survival constraints
 
        for (k in 1:npop) {
          for (t in first_pop[k]:(last_pop[k]-1)) {
            phi[1,k,t] <- ilogit(alpha[1] + beta*(density[k,t]-1) + epsilon[t])
            phi[2,k,t] <- ilogit(alpha[2] + beta*(density[k,t]-1) + epsilon[t])
          }#k
        }#t

      for (i in 1:nind) {

        # Part 2.2: Residency Process

        R[i] ~ dbern(pi[stage[i,first[i]]])
        r[i] ~ dbern(R[i]*rho[stage[i,first[i]]])

        # Part 2.3: Survival Process
        
        z[i,first[i]] ~ dbern(1)
      
        for (t in first[i]:(last_pop[pop[i]]-1)) {
          z[i,t+1] ~ dbern(phi[stage[i,t],pop[i],t]*z[i,t]*R[i])
        }#t
      
        # Part 2.4: Observation Process
        
        y[i,first[i],first_sub[i]] ~ dbern(1)
        y_pred[i,first[i],first_sub[i]] ~ dbern(1)
      
        for(h in (first_sub[i]+1):nsub) {
          p[i,first[i],h] <- ilogit(gamma[stage[i,first[i]]] + delta*effort[pop[i],first[i],h])
          y[i,first[i],h] ~ dbern(z[i,first[i]]*p[i,first[i],h])
          y_pred[i,first[i],h] ~ dbern(z[i,first[i]]*p[i,first[i],h])
        }#h
      
        for (t in first[i]:(last_pop[pop[i]]-1)) {
          for(h in 1:nsub) {
            p[i,t+1,h] <- ilogit(gamma[stage[i,t+1]] + delta*effort[pop[i],t+1,h])
            y[i,t+1,h] ~ dbern(z[i,t+1]*p[i,t+1,h])
            y_pred[i,t+1,h] ~ dbern(z[i,t+1]*p[i,t+1,h])
          }#h
        }#t
      }#i
      
      # Part 3: Population size and Density Estimation
      
        # Part 3.1: Probability of capture per population and year
        
        for (k in 1:npop) {
          for (t in first_pop[k]:last_pop[k]) {
            py_j[k,t] <- 1 - ((1-ilogit(gamma[1] + delta*effort[k,t,1]))*
                              (1-ilogit(gamma[1] + delta*effort[k,t,2]))*
                              (1-ilogit(gamma[1] + delta*effort[k,t,3]))*
                              (1-ilogit(gamma[1] + delta*effort[k,t,4])))
      
            py_a[k,t] <- 1 - ((1-ilogit(gamma[2] + delta*effort[k,t,1]))*
                              (1-ilogit(gamma[2] + delta*effort[k,t,2]))*
                              (1-ilogit(gamma[2] + delta*effort[k,t,3]))*
                              (1-ilogit(gamma[2] + delta*effort[k,t,4])))
          }#t
        }#k

        # Part 3.2: Density estimation
        
        for (k in 1:npop) {
          for (t in first_pop[k]:last_pop[k]) {
            Nad[k,t] <- Nobs_ad[k,t]/py_a[k,t] + (1-py_a[k,t])/py_a[k,t]
            Njuv1[k,t] <- Nobs_juv1[k,t]/py_j[k,t] + (1-py_j[k,t])/py_j[k,t]
      
            ## Total population size      
            N_temp[k,t] <- Nad[k,t] + Njuv1[k,t]
            N[k,t] <- ifelse(effort_year[k,t]==0, 0, N_temp[k,t])
      
            ## Count the number of years without effort in a population. 
            ## This is used for calculating mean pop size.
            n1[k,t] <- equals(effort_year[k,t], 0)
      
            ## Population Density (Assign 1 if there is no effort)
            density[k,t] <- ifelse(effort_year[k,t]==0, 1, N[k,t]/mean.N[k])
          }#t
      
          ## Counting the number of populations with effort across study period
          n2[k] <- (last_pop[k] - first_pop[k] + 1) - sum(n1[k,first_pop[k]:last_pop[k]])
      
          ## Mean pop size of effort years
          mean.N[k] <- sum(N[k,first_pop[k]:last_pop[k]])/n2[k]
        }#k
      
      # Part 4: Fecundity model
      
      for (l in 1:nfec) {
        year[l] <- year_index[l]-1
        density_fec[l] <- density[pop_index[l],year[l]]
        Njuv2[l] <- exp(log(Nad[pop_index[l],year_index[l]]) + theta + zeta*(density_fec[l]-1) + 
                        omega[l])
        w[l] ~ dbern(psi)
        Nobs_juv2[l] ~ dpois(max(py_j[pop_index[l],year_index[l]]*Njuv2[l]*w[l], 0.00000001))
        Nobs_juv_pred[l] ~ dpois(max(py_j[pop_index[l],year_index[l]]*Njuv2[l]*w[l], 0.00000001))
      }#l
      
      # Part 5: Model Checking
      
        # Part 5.1: Fecundity
      
        for (l in 1:nfec) {
          ft_fec_data1[l] <- (sqrt(Nobs_juv2[l])-sqrt(py_j[pop_index[l],year_index[l]]*Njuv2[l]*w[l]))^2
          ft_fec_pred1[l] <- (sqrt(Nobs_juv_pred[l])-sqrt(py_j[pop_index[l],year_index[l]]*Njuv2[l]*w[l]))^2
        }#l
      
        ft_fec_data2 <- sum(ft_fec_data1[])
        ft_fec_pred2 <- sum(ft_fec_pred1[])
      
        pvalue_fec <- step(ft_fec_pred2-ft_fec_data2)
      
        # Part 5.2: Capture History
      
        for (i in 1:nind) {
          ### The next two lines are adjustments for code to run
          ### They don't affect the calculation of Freeman-Tukey statistic
          p[i,first[i],first_sub[i]] <- 1
          start[i] <- ifelse(first_sub[i]==nsub, nsub, first_sub[i]+1)
      
          ts[i] <- (sqrt(sum(y[i,first[i],start[i]:nsub])) - 
                    sqrt(sum(p[i,first[i],start[i]:nsub])))^2
          ft1_data[i,first[i]] <- ifelse(first_sub[i]==nsub, 0, ts[i])
      
          ts_pred[i] <- (sqrt(sum(y_pred[i,first[i],start[i]:nsub])) - 
                         sqrt(sum(p[i,first[i],start[i]:nsub])))^2
          ft1_pred[i,first[i]] <- ifelse(first_sub[i]==nsub, 0, ts_pred[i])
      
          for (t in (first[i]+1):(last_pop[pop[i]])) {
            ft1_data[i,t] <- (sqrt(sum(y[i,t,])) - sqrt(z[i,t]*sum(p[i,t,])))^2
            ft1_pred[i,t] <- (sqrt(sum(y_pred[i,t,])) - sqrt(z[i,t]*sum(p[i,t,])))^2
          }#t
      
          ft2_data[i] <- sum(ft1_data[i,first[i]:last_pop[pop[i]]])
          ft2_pred[i] <- sum(ft1_pred[i,first[i]:last_pop[pop[i]]])
      }#i
      
      ft3_data <- sum(ft2_data[])
      ft3_pred <- sum(ft2_pred[])
      
      pvalue_ch <- step(ft3_pred-ft3_data)
      
      }", fill = TRUE)
  sink()  
}
```


### 2. Density Dependent CJS-pop without Residents

```{r, message=FALSE}

{
sink("cjspop_nores.jags")
  cat("
      model {
      
      # Part 1: Priors
      
        # Part 1.1: Priors for survival model

        survival_ad ~ dunif(0,1)
        survival_juv <- (1-survival_ad)/fecundity
        alpha[2] <- logit(survival_ad)
        alpha[1] <- logit(survival_juv)
        beta ~ dt(0,pow(10,-2),1)
        sigma_s ~ dt(0,pow(10,-2),1)T(0,)
        tau_s <- 1/(sigma_s^2)

        for (t in 1:(nyear-1)) {
          epsilon[t] ~ dnorm(0, tau_s)
        }

        # Part 1.2: Priors for capture porbablity model

        gamma[1] ~ dt(0,pow(10,-2),1)
        gamma[2] ~ dt(0,pow(10,-2),1)
        delta ~ dt(0,pow(2.5,-2),1)
      
        # Part 1.3: Priors for fecundity model

        fecundity ~ dnorm(0,pow(10,-2))T(0,)
        theta <- log(fecundity)
        zeta ~ dt(0,pow(5,-2),1)
        sigma_f ~ dt(0,pow(5,-2),1)T(0,)
        tau_f <- 1/(sigma_f^2)
        psi ~ dunif(0,1)

        for (l in 1:nfec) {
          omega[l] ~ dnorm(0, tau_f)
        }

      # Part 2: Robust design mark-recapture model

        # Part 2.1: Survival constraints
 
        for (k in 1:npop) {
          for (t in first_pop[k]:(last_pop[k]-1)) {
            phi[1,k,t] <- ilogit(alpha[1] + beta*(density[k,t]-1) + epsilon[t])
            phi[2,k,t] <- ilogit(alpha[2] + beta*(density[k,t]-1) + epsilon[t])
          }#k
        }#t

      for (i in 1:nind) {

        # Part 2.2: Survival Process
        
        z[i,first[i]] ~ dbern(1)
      
        for (t in first[i]:(last_pop[pop[i]]-1)) {
          z[i,t+1] ~ dbern(phi[stage[i,t],pop[i],t]*z[i,t])
        }#t
      
        # Part 2.4: Observation Process
        
        y[i,first[i],first_sub[i]] ~ dbern(1)
        y_pred[i,first[i],first_sub[i]] ~ dbern(1)
      
        for(h in (first_sub[i]+1):nsub) {
          p[i,first[i],h] <- ilogit(gamma[stage[i,first[i]]] + delta*effort[pop[i],first[i],h])
          y[i,first[i],h] ~ dbern(z[i,first[i]]*p[i,first[i],h])
          y_pred[i,first[i],h] ~ dbern(z[i,first[i]]*p[i,first[i],h])
        }#h
      
        for (t in first[i]:(last_pop[pop[i]]-1)) {
          for(h in 1:nsub) {
            p[i,t+1,h] <- ilogit(gamma[stage[i,t+1]] + delta*effort[pop[i],t+1,h])
            y[i,t+1,h] ~ dbern(z[i,t+1]*p[i,t+1,h])
            y_pred[i,t+1,h] ~ dbern(z[i,t+1]*p[i,t+1,h])
          }#h
        }#t
      }#i
      
      # Part 3: Population size and Density Estimation
      
        # Part 3.1: Probability of capture per population and year
        
        for (k in 1:npop) {
          for (t in first_pop[k]:last_pop[k]) {
            py_j[k,t] <- 1 - ((1-ilogit(gamma[1] + delta*effort[k,t,1]))*
                              (1-ilogit(gamma[1] + delta*effort[k,t,2]))*
                              (1-ilogit(gamma[1] + delta*effort[k,t,3]))*
                              (1-ilogit(gamma[1] + delta*effort[k,t,4])))
      
            py_a[k,t] <- 1 - ((1-ilogit(gamma[2] + delta*effort[k,t,1]))*
                              (1-ilogit(gamma[2] + delta*effort[k,t,2]))*
                              (1-ilogit(gamma[2] + delta*effort[k,t,3]))*
                              (1-ilogit(gamma[2] + delta*effort[k,t,4])))
          }#t
        }#k

        # Part 3.2: Density estimation
        
        for (k in 1:npop) {
          for (t in first_pop[k]:last_pop[k]) {
            Nad[k,t] <- Nobs_ad[k,t]/py_a[k,t] + (1-py_a[k,t])/py_a[k,t]
            Njuv1[k,t] <- Nobs_juv1[k,t]/py_j[k,t] + (1-py_j[k,t])/py_j[k,t]
      
            ## Total population size      
            N_temp[k,t] <- Nad[k,t] + Njuv1[k,t]
            N[k,t] <- ifelse(effort_year[k,t]==0, 0, N_temp[k,t])
      
            ## Count the number of years without effort in a population. 
            ## This is used for calculating mean pop size.
            n1[k,t] <- equals(effort_year[k,t], 0)
      
            ## Population Density (Assign 1 if there is no effort)
            density[k,t] <- ifelse(effort_year[k,t]==0, 1, N[k,t]/mean.N[k])
          }#t
      
          ## Counting the number of populations with effort across study period
          n2[k] <- (last_pop[k] - first_pop[k] + 1) - sum(n1[k,first_pop[k]:last_pop[k]])
      
          ## Mean pop size of effort years
          mean.N[k] <- sum(N[k,first_pop[k]:last_pop[k]])/n2[k]
        }#k
      
      # Part 4: Fecundity model
      
      for (l in 1:nfec) {
        year[l] <- year_index[l]-1
        density_fec[l] <- density[pop_index[l],year[l]]
        Njuv2[l] <- exp(log(Nad[pop_index[l],year_index[l]]) + theta + zeta*(density_fec[l]-1) + 
                        omega[l])
        w[l] ~ dbern(psi)
        Nobs_juv2[l] ~ dpois(max(py_j[pop_index[l],year_index[l]]*Njuv2[l]*w[l], 0.00000001))
        Nobs_juv_pred[l] ~ dpois(max(py_j[pop_index[l],year_index[l]]*Njuv2[l]*w[l], 0.00000001))
      }#l
      
      # Part 5: Model Checking
      
        # Part 5.1: Fecundity
      
        for (l in 1:nfec) {
          ft_fec_data1[l] <- (sqrt(Nobs_juv2[l])-sqrt(py_j[pop_index[l],year_index[l]]*Njuv2[l]*w[l]))^2
          ft_fec_pred1[l] <- (sqrt(Nobs_juv_pred[l])-sqrt(py_j[pop_index[l],year_index[l]]*Njuv2[l]*w[l]))^2
        }#l
      
        ft_fec_data2 <- sum(ft_fec_data1[])
        ft_fec_pred2 <- sum(ft_fec_pred1[])
      
        pvalue_fec <- step(ft_fec_pred2-ft_fec_data2)
      
        # Part 5.2: Capture History
      
        for (i in 1:nind) {
          ### The next two lines are adjustments for code to run
          ### They don't affect the calculation of Freeman-Tukey statistic
          p[i,first[i],first_sub[i]] <- 1
          start[i] <- ifelse(first_sub[i]==nsub, nsub, first_sub[i]+1)
      
          ts[i] <- (sqrt(sum(y[i,first[i],start[i]:nsub])) - 
                    sqrt(sum(p[i,first[i],start[i]:nsub])))^2
          ft1_data[i,first[i]] <- ifelse(first_sub[i]==nsub, 0, ts[i])
      
          ts_pred[i] <- (sqrt(sum(y_pred[i,first[i],start[i]:nsub])) - 
                         sqrt(sum(p[i,first[i],start[i]:nsub])))^2
          ft1_pred[i,first[i]] <- ifelse(first_sub[i]==nsub, 0, ts_pred[i])
      
          for (t in (first[i]+1):(last_pop[pop[i]])) {
            ft1_data[i,t] <- (sqrt(sum(y[i,t,])) - sqrt(z[i,t]*sum(p[i,t,])))^2
            ft1_pred[i,t] <- (sqrt(sum(y_pred[i,t,])) - sqrt(z[i,t]*sum(p[i,t,])))^2
          }#t
      
          ft2_data[i] <- sum(ft1_data[i,first[i]:last_pop[pop[i]]])
          ft2_pred[i] <- sum(ft1_pred[i,first[i]:last_pop[pop[i]]])
      }#i
      
      ft3_data <- sum(ft2_data[])
      ft3_pred <- sum(ft2_pred[])
      
      pvalue_ch <- step(ft3_pred-ft3_data)
      
      }", fill = TRUE)
  sink()  
}
```


### 3. Density Independent CJS-pop with Residents

```{r, message=FALSE}

{
sink("cjspop_di.jags")
  cat("
      model {
      
      # Part 1: Priors
      
        # Part 1.1: Priors for survival model

        survival_ad ~ dunif(0,1)
        survival_juv <- (1-survival_ad)/fecundity
        alpha[2] <- logit(survival_ad)
        alpha[1] <- logit(survival_juv)
        sigma_s ~ dt(0,pow(10,-2),1)T(0,)
        tau_s <- 1/(sigma_s^2)

        for (t in 1:(nyear-1)) {
          epsilon[t] ~ dnorm(0, tau_s)
        }

        # Part 1.2: Priors for capture porbablity model

        gamma[1] ~ dt(0,pow(10,-2),1)
        gamma[2] ~ dt(0,pow(10,-2),1)
        delta ~ dt(0,pow(2.5,-2),1)
      
        # Part 1.3: Priors for fecundity model

        fecundity ~ dnorm(0,pow(10,-2))T(0,)
        theta <- log(fecundity)
        sigma_f ~ dt(0,pow(5,-2),1)T(0,)
        tau_f <- 1/(sigma_f^2)
        psi ~ dunif(0,1)

        for (l in 1:nfec) {
          omega[l] ~ dnorm(0, tau_f)
        }
        
        # Part 1.4: Priors for residency model
      
        pi[1] ~ dunif(0,1)
        pi[2] ~ dunif(0,1)
        rho[1] ~ dunif(0,1)
        rho[2] ~ dunif(0,1)

      # Part 2: Robust design mark-recapture model

      for (i in 1:nind) {

        # Part 2.1: Residency Process

        R[i] ~ dbern(pi[stage[i,first[i]]])
        r[i] ~ dbern(R[i]*rho[stage[i,first[i]]])

        # Part 2.2: Survival Process
        
        z[i,first[i]] ~ dbern(1)
      
        for (t in first[i]:(last_pop[pop[i]]-1)) {
          phi[i,t] <- ilogit(alpha[stage[i,t]] + epsilon[t])
          z[i,t+1] ~ dbern(phi[i,t]*z[i,t]*R[i])
        }#t
      
        # Part 2.3: Observation Process
        
        y[i,first[i],first_sub[i]] ~ dbern(1)
        y_pred[i,first[i],first_sub[i]] ~ dbern(1)
      
        for(h in (first_sub[i]+1):nsub) {
          p[i,first[i],h] <- ilogit(gamma[stage[i,first[i]]] + delta*effort[pop[i],first[i],h])
          y[i,first[i],h] ~ dbern(z[i,first[i]]*p[i,first[i],h])
          y_pred[i,first[i],h] ~ dbern(z[i,first[i]]*p[i,first[i],h])
        }#h
      
        for (t in first[i]:(last_pop[pop[i]]-1)) {
          for(h in 1:nsub) {
            p[i,t+1,h] <- ilogit(gamma[stage[i,t+1]] + delta*effort[pop[i],t+1,h])
            y[i,t+1,h] ~ dbern(z[i,t+1]*p[i,t+1,h])
            y_pred[i,t+1,h] ~ dbern(z[i,t+1]*p[i,t+1,h])
          }#h
        }#t
      }#i
      
      # Part 3: Population size and Density Estimation
      
        # Part 3.1: Probability of capture per population and year
        
        for (k in 1:npop) {
          for (t in first_pop[k]:last_pop[k]) {
            py_j[k,t] <- 1 - ((1-ilogit(gamma[1] + delta*effort[k,t,1]))*
                              (1-ilogit(gamma[1] + delta*effort[k,t,2]))*
                              (1-ilogit(gamma[1] + delta*effort[k,t,3]))*
                              (1-ilogit(gamma[1] + delta*effort[k,t,4])))

            py_a[k,t] <- 1 - ((1-ilogit(gamma[2] + delta*effort[k,t,1]))*
                              (1-ilogit(gamma[2] + delta*effort[k,t,2]))*
                              (1-ilogit(gamma[2] + delta*effort[k,t,3]))*
                              (1-ilogit(gamma[2] + delta*effort[k,t,4])))
          }#t
        }#k

        # Part 3.2: Adult Population Size Estimation
        
        for (k in 1:npop) {
          for (t in first_pop[k]:last_pop[k]) {
            Nad[k,t] <- Nobs_ad[k,t]/py_a[k,t] + (1-py_a[k,t])/py_a[k,t]
          }#t
        }#k
      
      # Part 4: Fecundity model
      
      for (l in 1:nfec) {
        Njuv2[l] <- exp(log(Nad[pop_index[l],year_index[l]]) + theta + omega[l])
        w[l] ~ dbern(psi)
        Nobs_juv2[l] ~ dpois(max(py_j[pop_index[l],year_index[l]]*Njuv2[l]*w[l], 0.00000001))
        Nobs_juv_pred[l] ~ dpois(max(py_j[pop_index[l],year_index[l]]*Njuv2[l]*w[l], 0.00000001))
      }#l
      
      # Part 5: Model Checking
      
        # Part 5.1: Fecundity
      
        for (l in 1:nfec) {
          ft_fec_data1[l] <- (sqrt(Nobs_juv2[l])-sqrt(py_j[pop_index[l],year_index[l]]*Njuv2[l]*w[l]))^2
          ft_fec_pred1[l] <- (sqrt(Nobs_juv_pred[l])-sqrt(py_j[pop_index[l],year_index[l]]*Njuv2[l]*w[l]))^2
        }#l
      
        ft_fec_data2 <- sum(ft_fec_data1[])
        ft_fec_pred2 <- sum(ft_fec_pred1[])
      
        pvalue_fec <- step(ft_fec_pred2-ft_fec_data2)
      
        # Part 5.2: Capture History
      
        for (i in 1:nind) {
          ### The next two lines are adjustments for code to run
          ### They don't affect the calculation of Freeman-Tukey statistic
          p[i,first[i],first_sub[i]] <- 1
          start[i] <- ifelse(first_sub[i]==nsub, nsub, first_sub[i]+1)
      
          ts[i] <- (sqrt(sum(y[i,first[i],start[i]:nsub])) - 
                    sqrt(sum(p[i,first[i],start[i]:nsub])))^2
          ft1_data[i,first[i]] <- ifelse(first_sub[i]==nsub, 0, ts[i])
      
          ts_pred[i] <- (sqrt(sum(y_pred[i,first[i],start[i]:nsub])) - 
                         sqrt(sum(p[i,first[i],start[i]:nsub])))^2
          ft1_pred[i,first[i]] <- ifelse(first_sub[i]==nsub, 0, ts_pred[i])
      
          for (t in (first[i]+1):(last_pop[pop[i]])) {
            ft1_data[i,t] <- (sqrt(sum(y[i,t,])) - sqrt(z[i,t]*sum(p[i,t,])))^2
            ft1_pred[i,t] <- (sqrt(sum(y_pred[i,t,])) - sqrt(z[i,t]*sum(p[i,t,])))^2
          }#t
      
          ft2_data[i] <- sum(ft1_data[i,first[i]:last_pop[pop[i]]])
          ft2_pred[i] <- sum(ft1_pred[i,first[i]:last_pop[pop[i]]])
      }#i
      
      ft3_data <- sum(ft2_data[])
      ft3_pred <- sum(ft2_pred[])
      
      pvalue_ch <- step(ft3_pred-ft3_data)
      
      }", fill = TRUE)
  sink()  
}

```


### 4. Simple CJS with Residents

```{r, message=FALSE}

{
sink("cjs_r.jags")
  cat("
      model {
      
      # Part 1: Priors
      
        # Part 1.1: Priors for survival model

        survival_ad ~ dunif(0,1)
        survival_juv ~ dunif(0,1)
        alpha[2] <- logit(survival_ad)
        alpha[1] <- logit(survival_juv)
        sigma_s ~ dt(0,pow(10,-2),1)T(0,)
        tau_s <- 1/(sigma_s^2)

        for (t in 1:(nyear-1)) {
          epsilon[t] ~ dnorm(0, tau_s)
        }

        # Part 1.2: Priors for capture porbablity model

        gamma[1] ~ dt(0,pow(10,-2),1)
        gamma[2] ~ dt(0,pow(10,-2),1)
        delta ~ dt(0,pow(2.5,-2),1)
        
        # Part 1.3: Priors for residency model
      
        pi[1] ~ dunif(0,1)
        pi[2] ~ dunif(0,1)
        rho[1] ~ dunif(0,1)
        rho[2] ~ dunif(0,1)

      # Part 2: Robust design mark-recapture model

      for (i in 1:nind) {

        # Part 2.1: Residency Process

        R[i] ~ dbern(pi[stage[i,first[i]]])
        r[i] ~ dbern(R[i]*rho[stage[i,first[i]]])

        # Part 2.2: Survival Process
        
        z[i,first[i]] ~ dbern(1)
      
        for (t in first[i]:(last_pop[pop[i]]-1)) {
          phi[i,t] <- ilogit(alpha[stage[i,t]] + epsilon[t])
          z[i,t+1] ~ dbern(phi[i,t]*z[i,t]*R[i])
        }#t
      
        # Part 2.3: Observation Process
        
        y[i,first[i],first_sub[i]] ~ dbern(1)
        y_pred[i,first[i],first_sub[i]] ~ dbern(1)
      
        for(h in (first_sub[i]+1):nsub) {
          p[i,first[i],h] <- ilogit(gamma[stage[i,first[i]]] + delta*effort[pop[i],first[i],h])
          y[i,first[i],h] ~ dbern(z[i,first[i]]*p[i,first[i],h])
          y_pred[i,first[i],h] ~ dbern(z[i,first[i]]*p[i,first[i],h])
        }#h
      
        for (t in first[i]:(last_pop[pop[i]]-1)) {
          for(h in 1:nsub) {
            p[i,t+1,h] <- ilogit(gamma[stage[i,t+1]] + delta*effort[pop[i],t+1,h])
            y[i,t+1,h] ~ dbern(z[i,t+1]*p[i,t+1,h])
            y_pred[i,t+1,h] ~ dbern(z[i,t+1]*p[i,t+1,h])
          }#h
        }#t
      }#i
      
      # Part 3: Model Checking
      
        for (i in 1:nind) {
          ### The next two lines are adjustments for code to run
          ### They don't affect the calculation of Freeman-Tukey statistic
          p[i,first[i],first_sub[i]] <- 1
          start[i] <- ifelse(first_sub[i]==nsub, nsub, first_sub[i]+1)
      
          ts[i] <- (sqrt(sum(y[i,first[i],start[i]:nsub])) - 
                    sqrt(sum(p[i,first[i],start[i]:nsub])))^2
          ft1_data[i,first[i]] <- ifelse(first_sub[i]==nsub, 0, ts[i])
      
          ts_pred[i] <- (sqrt(sum(y_pred[i,first[i],start[i]:nsub])) - 
                         sqrt(sum(p[i,first[i],start[i]:nsub])))^2
          ft1_pred[i,first[i]] <- ifelse(first_sub[i]==nsub, 0, ts_pred[i])
      
          for (t in (first[i]+1):(last_pop[pop[i]])) {
            ft1_data[i,t] <- (sqrt(sum(y[i,t,])) - sqrt(z[i,t]*sum(p[i,t,])))^2
            ft1_pred[i,t] <- (sqrt(sum(y_pred[i,t,])) - sqrt(z[i,t]*sum(p[i,t,])))^2
          }#t
      
          ft2_data[i] <- sum(ft1_data[i,first[i]:last_pop[pop[i]]])
          ft2_pred[i] <- sum(ft1_pred[i,first[i]:last_pop[pop[i]]])
      }#i
      
      ft3_data <- sum(ft2_data[])
      ft3_pred <- sum(ft2_pred[])
      
      pvalue_ch <- step(ft3_pred-ft3_data)
      
      }", fill = TRUE)
  sink()  
}
```
````
