## Appendix S3-S4 for "Estimating fecundity and density dependence from mark-recapture data for making population projections": AppendixS4.Rmd

````
---
title: "Data Generation"
output:
  html_document:
    df_print: paged
---

### Introduction

Density dependent and stochastic population simulations used for generating capture history data. I first introduce the functions to simulate population dynamics and to extract mark-recapture data. Then I generate data under different scenarios.


#### Load packages

```{r}
library(abind)
library(boot)
library(tidyverse)

```


### 1. Population and capture simulation functions

This is a stage matrix model with two stages. However, I can't run this with typical matrix math because I have to follow the fate of each individual in the population. So I have to go through each individual in each population in each time step and decide if that individual survived in that time step. This is the indvidual-level component of the simulation. The total population size divided by carrying capacity and along with a random and temporal variance component determines the survival rate in each year and each population. This is the population-level compoenent of the simulation, and also the survival rate I use to determine individual survival.

Fecundity is modelled in simpler terms. It doesn't matter for the simulation's sake which individual bred how much. So fecundity component is strictly a population-level model. Number of newborn individuals each year in each population comes from a Poisson distribtuion whose mean is calculated by multiplying fecundity with number of adults in that population. Fecundity, like survival, is a function of density and random temporal variation.


#### Function 1: survive

**z**: State matrix

**t**: Time step 

**s**: Survival rate 

z is the state matrix with number of rows equal to number of individuals and number columns equal to number of time steps. 1 indicates an indivial (row) is alive in a given time step (column), and 0 indicates that individual is dead. Function loops through all individuals and decides their state on time step t+1. The state in time t+1 is condiontional on the state in time t. Survival is determined as Bernoulli draw with probablity equal to survival rate. 

```{r}
survive <- function(z, t, s) {
  for (i in 1:nrow(z)) {
    z[i,t+1] <- z[i,t]*rbinom(1, 1, s)
  }#i
  return(z)
}
```


#### Function 2: setup.sim

**K**: Carrying capacity. Assumed to be the same across populations.

**npop**: Number of populations in the simulation.

**time**: Number of time steps to run the simulation.

Sets up the starting values for the population simulation. Notal, Nad, and Njuv are total, adult and juvenile population sizes. First year values are filled assuming each is half the carrying capacity and population in total is at carrying capacity. z.ad and z.juv are state matrices for adults and juveniles. Each population have their own state matrix. All the starting individuals are assumed to be alive in the first time step. stage is the matrix with the stage information for each individual(1:Adult, 2:Juvenile). Each population has it's own stage matrix. Only adults are put in the stage matrix at first. Juveniles are added as simulation is run, as can be seen below in the **simulate.state** function.

```{r}
setup.sim <- function(K, npop, time) {
  Ntotal <- matrix(nrow = npop, ncol = time)
  Nad <- matrix(nrow = npop, ncol = time)
  Njuv <- matrix(nrow = npop, ncol = time)
  
  # First year population size
  K <- rep(K, npop)
  Nad[,1] <- K/2
  Njuv[,1] <- K/2
  Ntotal[,1] <- Nad[,1] + Njuv[,1]
  
  # State matrix
  z.ad <- split(Nad[,1], 1:npop)
  z.ad <- lapply(z.ad, function(x) matrix(ncol = time, nrow = x))
  z.juv <- split(Njuv[,1], 1:npop)
  z.juv <- lapply(z.juv, function(x) matrix(ncol = time, nrow = x))
  
  # Stage matrix
  stage <- split(Nad[,1], 1:npop)
  stage <- lapply(stage, function(x) matrix(ncol = time, nrow = x))
  
  # First year state
  for (p in 1:npop) {
    z.ad[[p]][,1] <- 1
    z.juv[[p]][,1] <- 1
    stage[[p]][,1:time] <- 2
  }
  
  results <- list(Nad = Nad,
                  Njuv = Njuv,
                  Ntotal = Ntotal,
                  K = K,
                  z.ad = z.ad,
                  z.juv = z.juv,
                  stage = stage,
                  time = time,
                  npop = npop,
                  K = K)
  
  return(results)
}
```


#### Function 3: simulate.state

**sv**: starting values setup with **setup.sim** function.

**alpha1**: Juvenile survival in logit scale when density is 0.

**alpha2**: Adult survival in logit scale when density is 0.

**beta**: Density dependence strength of survival in logit scale.

**zeta**: Density dependence strenth of fecundity in log scale.

**sigma_s**: Temporal process standard deviation of survival in logit scale.

**sigma_f**: Temporal process standard deviation of fecundity in log scale.

Simulates the state of each individual in each population at each time step, along with population size. Survival and fecundity is updated in each population at each time step as a function of population density and random temporal variance. **survival** function is used to update the state matrices. A new state matrix for juveniles (z.juv) is created in each population at each time step with newborns. After their survival to t+1 these individuals become adults and juvenile state matrix is added to adult state matrix (z.ad). That way adult state matrix in each population grows by row at each time step, while juvenile state matrix is "reset" with newborns. Similar logic applies to the stage matrix. Newborns are immediatly assigned 2 as juveniles, and all following years (irrespective of their survival) are assinged 1 as adults. This juvenile stage matrix then is added to the adult stage matrix.

Fecundity at 0 density (**theta**) is calculated using **alpha1**, **alpha2**, **beta**, and **zeta**. In a density dependent model lambda must be 1 at carrying capacity. If 4 parameters are known the 5th is not free to vary if the population is to be regulated around stated K. We use this relationship between vital rates to calculate fecundity.  

The general flow of the simulation can be followed like this
1- Survival rate of adults and juveniles are calculated in each population as a function of density at time t and random temporal variance.
2- Adult individuals survive to t+1.
3- If there are juveniles in a population at time t they survive to t+1 and become adults.
4- Adult population size at t+1 is calculated by summing up individuals alive at t+1.
5- Fecundity at t+1 is calculated as a function of density at time t and random temporal variance.
6- Adults at t+1 breed (Nad*f) and produce juveniles of t+1.
7- Total population size and density at t+1 is calculated by summing adults and juveniles and dividing by carrying capacity.
8- Iteration ends, loop goes back to step 1. This loop is repeated for each population at each time step.

```{r}
simulate.state <- function(sv, alpha1, alpha2, beta, zeta, sigma_s, sigma_f) {
  # Parameters
  time <- sv$time
  npop <- sv$npop
  K <- sv$K
  Ntotal <- sv$Ntotal
  Nad <- sv$Nad
  Njuv <- sv$Njuv
  z.ad <- sv$z.ad
  z.juv <- sv$z.juv
  stage <- sv$stage
  z <- list()
  
  s.ad <- matrix(nrow = npop, ncol = time)
  s.juv <- matrix(nrow = npop, ncol = time)
  f <- matrix(nrow = npop, ncol = time)
  theta <- log((1-(inv.logit(alpha2 + beta)))/(inv.logit(alpha1 + beta))) - zeta
  
  # Main loop
  for (t in 1:(time-1)) {
    
    for (p in 1:npop) {
      
      # random spatio-temporal variance component
      epsilon <- rnorm(1, 0, sigma_s)
      omega <- rnorm(1, 0, sigma_f)
      
      s.ad[p,t] <- inv.logit(alpha2 + beta*Ntotal[p,t]/K[p] + epsilon)
      s.juv[p,t] <- inv.logit(alpha1 + beta*Ntotal[p,t]/K[p] + epsilon)
      
      # Survival
      z.ad[[p]] <- survive(z.ad[[p]], t, s.ad[p,t])
      if (Njuv[p,t]>0) {
        # Juvenile stage
        stage.juv <- matrix(nrow = Njuv[p,t], ncol = time)
        stage.juv[,t] <- 1
        stage.juv[,(t+1):time] <- 2
        stage[[p]] <- rbind(stage[[p]], stage.juv)
        
        # Juvenile survival
        z.juv[[p]] <- survive(z.juv[[p]], t, s.juv[p,t])
        z.ad[[p]] <- rbind(z.ad[[p]], z.juv[[p]])
      } 
      
      # Fecundity
      f[p,t+1] <- exp(theta + zeta*Ntotal[p,t]/K[p] + omega)
      Nad[p,t+1] <- sum(z.ad[[p]][,t+1])
      Njuv[p,t+1] <- rpois(1, f[p,t+1]*Nad[p,t+1])
      Ntotal[p,t+1] <- Nad[p,t+1] + Njuv[p,t+1]
      
      # Empty juvenile matrix for t+1
      z.juv[[p]] <- matrix(nrow = Njuv[p,t+1], ncol = time)
      z.juv[[p]][,t+1] <- 1
      
      # Last year
      if ((t+1)==time) {
        if (Njuv[p,t+1]>0) {
          stage.juv <- matrix(nrow = Njuv[p,t+1], ncol = time)
          stage.juv[,t+1] <- 1
          stage[[p]] <- rbind(stage[[p]], stage.juv)
          z[[p]] <- rbind(z.ad[[p]], z.juv[[p]])
        } else z[[p]] <- z.ad[[p]]
      }
    }#p
  }#t
  
  parameters <- list(alpha1 = alpha1,
                     alpha2 = alpha2,
                     theta = theta,
                     beta = beta,
                     zeta = zeta,
                     sigma_s = sigma_s,
                     sigma_f = sigma_f)
  
  state_data <- list(Nad = Nad,
                     Njuv = Njuv,
                     Ntotal = Ntotal,
                     K = K, 
                     z = z, 
                     stage = stage,
                     s.ad = s.ad,
                     s.juv = s.juv,
                     f = f,
                     parameters = parameters)
  return(state_data)
}
```


#### Function 4: simulate.ch

**state**: Object created with **simulate.state** function.

**p.ad**: Adult capture probability in a sub-occasion.

**p.juv**: Juvenile capture probability in a sub-occasion

Generates a robust design capture history from the state matrix simulated with **simulate.state** function. Time steps in the population simulation above correspond to the primary occasions. Individuals can die between primary occasions (as simulated above). There are 4 sub-occassions within each primary occasion. Individuals do not die between sub-occasions, but they can be captured in each sub-occasion. This capture is conditional on them being alive in a given primary occasion. Capturing individuals is a Bernoulli draw with adult or juvenile capture probablity.    

```{r}
simulate.ch <- function(state, p.ad, p.juv) {
  z <- state$z
  stage <- state$stage
  nind <- sapply(z, nrow)
  npop <- length(state$K)
  time <- ncol(state$Ntotal)
  ch <- split(nind, 1:npop)
  ch <- lapply(ch, function(x) array(dim = c(x, time, 4)))
  
  for (p in 1:npop) {
    for (i in 1:nind[p]) {
      for (t in 1:time) {
        
        if (is.na(z[[p]][i,t])) {
          next
        } else {
          
          for (h in 1:4) {
              
              if (stage[[p]][i,t]==1) ch[[p]][i,t,h] <- rbinom(1,1,p.juv)*z[[p]][i,t]
              if (stage[[p]][i,t]==2) ch[[p]][i,t,h] <- rbinom(1,1,p.ad)*z[[p]][i,t]
          }
        }
      }
    }
  }
  return(ch)
}
```


### 2. Data formatting functions

Following functions format the data for Bayesian mark-recapture models in jags language.


#### Function 5: format.ch

**ch**: Capture history object generated with **simulate.ch**.

**state**: Object created with **simulate.state** function.

Removes individuals who are never captured from capture history, state, and stage matrices. Also creates a capture history matrix with only primary occasions.

```{r}
format.ch <- function(ch, state) {
  ch_robust <- ch
  stage <- state$stage
  npop <- length(ch_robust)
  
  
  # Removing individuals who were never captured
  mark_ind <- function(x) apply(x, 1, function(y) all(y==0, na.rm = T))
  index <- lapply(ch_robust, mark_ind)
                    
  for (p in 1:npop) {
    ch_robust[[p]] <- ch[[p]][!index[[p]],,]
    stage[[p]] <- stage[[p]][!index[[p]],]
  }
  
  # Collapse capture history into a yearly format
  collapse_sub <- function(x) apply(x, c(1,2), function(y) ifelse(any(y==1, na.rm = T), 1, 0))
  ch_year <- lapply(ch_robust, collapse_sub)
  
  results <- list(ch_robust = ch_robust,
                  ch_year = ch_year,
                  stage = stage)
  
  return(results)
}

```


#### Function 6: format.Nobs

**ch**: Object created with **format.ch** function.

Creates matrices for total number of captured adults and juveniles for all populations and time steps.

```{r}
format.Nobs <- function(ch) {
  ch_year <- ch$ch_year
  npop <- length(ch_year)
  stage <- ch$stage
  
  # Seperate adult and juvenile ch
  ch_ad <- list()
  ch_juv <- list()
  for (p in 1:npop) {
    ch_ad[[p]] <- ch_year[[p]]*ifelse(stage[[p]]==2, 1, 0)
    ch_juv[[p]] <- ch_year[[p]]*ifelse(stage[[p]]==1, 1, 0)
  }
  
  # Observed population sizes
  Nobs_ad <- lapply(ch_ad, function(x) apply(x, 2, sum, na.rm=T))
  Nobs_juv <- lapply(ch_juv, function(x) apply(x, 2, sum, na.rm=T))
  
  # Combine data across populations
  Nobs_ad <- do.call(rbind, Nobs_ad)
  Nobs_juv <- do.call(rbind, Nobs_juv)
  
  results <- list(Nobs_ad = Nobs_ad,
                  Nobs_juv = Nobs_juv,
                  npop = nrow(Nobs_ad))
  return(results)
}
```


#### Function 7: format.index

**ch**: Object created with **format.ch** function.

Creates multiple indices used in mark-recapture models in jags language.
1. First capture time step (prmiary occasion) of every individual.
2. First capture sub-occasion of every individual.
3. First capture time step of a population, in other words first time ever an individual was captured in a population.
4. Last capture time step of every individual.
5. Last capture time step of a population, in other words last time an individual was ever captured in a population.
6. Population number each individual belongs to.

```{r}
format.index <- function(ch) {
  ch_robust <- ch$ch_robust
  ch_year <- ch$ch_year
  npop <- length(ch_year)
  
  # First primary occasion capture
  detect_first <- function(x) apply(x, 1, function(x) min(which(x==1)))
  first <- lapply(ch_year, detect_first)
  
  # First sub-ccasion capture
  detect_first_sub <- function(x) {
    f <- min(which(x==1, arr.ind = T)[,1])
    return(min(which(x[f,]==1)))
  } 
  first_sub <- lapply(ch_robust, function(x) apply(x, 1, detect_first_sub))
  
  # First primary capture occasion of population
  first_pop <- lapply(ch_year, function(x) min(which(x==1, arr.in = T)[,2]))
  
  # Last primary capture occasion
  detect_last <- function(x) apply(x, 1, function(x) max(which(x==1)))
  last <- lapply(ch_year, detect_last)
  
  # Last primary capture occasion in a population
  last_pop <- lapply(ch_year, function(x) max(which(x==1, arr.in = T)[,2]))
  
  # Population index
  pop.list <- list()
  for (p in 1:npop) {
    pop.list[[p]] <- rep(p, times = nrow(ch_year[[p]]))
  }
  
  # Combine data across populations
  first <- do.call(c, first)
  first_sub <- do.call(c, first_sub)
  first_pop <- do.call(c, first_pop)
  last_pop <- do.call(c, last_pop)
  last <- do.call(c, last)
  pop <- do.call(c, pop.list)
  
  results <- list(first = first,
                  first_sub = first_sub,
                  first_pop = first_pop,
                  last = last,
                  last_pop = last_pop,
                  pop = pop)
}
```


#### Function 8: format.fec

**Nobs**: Object created with **format.Nobs** function.

Creates data for fecundity analysis. Filters data to populations and years with at least 1 adult and 1 juvenile capture. Also removes first year of capture in every population. That is because fecundity models use density from the previous year, and no such information is available for first year data.

```{r}
format.fec <- function(Nobs, index) {
  Nobs_ad <- Nobs$Nobs_ad
  Nobs_juv <- Nobs$Nobs_juv
  npop <- nrow(Nobs_ad)
  first_pop <- index$first_pop
  
  # Assign 0 population size to first capture year 
  for (i in 1:npop) {
    Nobs_ad[i,first_pop[i]] <- 0
    Nobs_juv[i,first_pop[i]] <- 0
  }
  
  # Population and year indices for fecundity data
  pop_index <- which(Nobs_ad>0 & Nobs_juv>0, arr.ind = T)[,1]
  year_index <- which(Nobs_ad>0 & Nobs_juv>0, arr.ind = T)[,2]
  
  # Fecundity data used in mark-recapture models.
  fec_data <- Nobs_juv[which(Nobs_ad>0 & Nobs_juv>0)]
  
  results <- list(fec_data = fec_data,
                  pop_index = pop_index,
                  year_index = year_index,
                  nfec = length(fec_data))
  
  return(results)
}
```


#### Function 9: wrap.chdata

**ch**: Capture history object generated with **simulate.ch**.

**state**: Object created with **simulate.state** function.

A wrapper function for functions 5 to 8. Formats the data to it's final form for mark-recapture analysis. Latent_state object is used to fasten the bayesian mark-recapture models in jags language. It sets the latent state of every individual to 1 between their first and last capture events. Their first year capture and time steps after the last capture year is set to NA. This way model does not try to estimate the state of individuals between their first and last capture years, which makes sense because we know they are alive in that time period. 

```{r}
wrap.chdata <- function(ch, state) {
  ch_mark <- format.ch(ch, state)
  Nobs <- format.Nobs(ch_mark)
  index <- format.index(ch_mark)
  fec <- format.fec(Nobs, index)
  
  # Combine data from populations
  ch_robust <- do.call(abind, list(ch_mark$ch_robust, along = 1))
  ch_year <- do.call(rbind, ch_mark$ch_year)
  stage <- do.call(rbind, ch_mark$stage)
  for (i in 1:nrow(ch_year)) {
    if (index$first[i]==1) next
    ch_year[i,1:(index$first[i]-1)] <- NA
  }
  
  # Latent state set up
  latent_state <- ch_year  
  for (i in 1:nrow(ch_robust)) {
    n1 <- min(which(ch_year[i,]==1))
    n2 <- max(which(ch_year[i,]==1))
    latent_state[i, n1:n2] <- 1
    latent_state[i, n1] <- NA 
  }
  latent_state[latent_state==0] <- NA
  
  results <- list(ch = ch_robust,
                  ch_year = ch_year,
                  stage = stage,
                  Nobs_ad = Nobs$Nobs_ad,
                  Nobs_juv = Nobs$Nobs_juv,
                  first = index$first,
                  first_pop = index$first_pop,
                  first_sub = index$first_sub,
                  last_pop = index$last_pop,
                  last = index$last,
                  pop = index$pop,
                  pop_index = fec$pop_index,
                  year_index = fec$year_index,
                  fec_data = fec$fec_data,
                  latent_state = latent_state,
                  nind = nrow(ch_year),
                  nyear = ncol(ch_year),
                  npop = Nobs$npop,
                  nfec = fec$nfec)
  return(results)
}
```


### 3. Data Generation

```{r}
n_iter <- 56
pop <- c(1,5,10)
K <- c(50,100,150)
dd <- c(-0.05, -0.5, -1)
dd_name <- c("l", "m", "h")

for (j in 1:length(pop)) {
  
  for (h in 1:length(K)) {
    
    for (d in 1:length(dd)) {
      sv <- setup.sim(npop = pop[j], time = 17, K = K[h])
    
      state <- list()
      ch <- list()
      chdata <- list()
    
      for (i in 1:n_iter) {
        state[[i]] <- simulate.state(sv = sv,
                                     alpha2 = 1,
                                     alpha1 = 0.5,
                                     beta = dd[d],
                                     zeta = dd[d],
                                     sigma_s = 0.5,
                                     sigma_f = 0.5)

        ch[[i]] <- simulate.ch(state[[i]],
                               p.ad = 0.10,
                               p.juv = 0.02)

        chdata[[i]] <- wrap.chdata(ch[[i]], state[[i]])
      }
      
      state_file <- paste("state", dd_name[d], sep = "_") %>%
        paste("p", sep = "_") %>%
        paste0(pop[j]) %>%
        paste("k", sep = "_") %>%
        paste0(K[h]) %>%
        paste("rds", sep = ".")
      
      ch_file <- paste("ch", dd_name[d], sep = "_") %>%
        paste("p", sep = "_") %>%
        paste0(pop[j]) %>%
        paste("k", sep = "_") %>%
        paste0(K[h]) %>%
        paste("rds", sep = ".")
      
      chdata_file <- paste("chdata", dd_name[d], sep = "_") %>%
        paste("p", sep = "_") %>%
        paste0(pop[j]) %>%
        paste("k", sep = "_") %>%
        paste0(K[h]) %>%
        paste("rds", sep = ".")
      
      saveRDS(state, file = paste("data", state_file, sep = "/"))
      saveRDS(ch, file = paste("data", ch_file, sep = "/"))
      saveRDS(chdata, file = paste("data", chdata_file, "sep = /"))
    }
  }
}
```

#### Simulation example data

```{r}

sv <- setup.sim(npop = 10, time = 17, K = 150)
    
state <- list()
ch <- list()
chdata <- list()

for (i in 1:2) {
  
        state[[i]] <- simulate.state(sv = sv,
                                     alpha2 = 1,
                                     alpha1 = 0.5,
                                     beta = -0.5,
                                     zeta = -0.5,
                                     sigma_s = 0.5,
                                     sigma_f = 0.5)

        ch[[i]] <- simulate.ch(state[[i]],
                               p.ad = 0.10,
                               p.juv = 0.02)

        chdata[[i]] <- wrap.chdata(ch[[i]], state[[i]])
}

saveRDS(chdata, file = "chdata_sim_example.rds")

```
````
